# High affinity chimeric antigen receptor signaling induces an inflammatory program in human regulatory T cells

**DOI:** 10.1101/2024.03.31.587467

**Authors:** Russell W. Cochrane, Rob A. Robino, Bryan Granger, Eva Allen, Silvia Vaena, Martin J. Romeo, Aguirre A. de Cubas, Stefano Berto, Leonardo M.R. Ferreira

## Abstract

Regulatory T cells (Tregs) are promising cellular therapies to induce immune tolerance in organ transplantation and autoimmune disease. The success of chimeric antigen receptor (CAR) T-cell therapy for cancer has sparked interest in using CARs to generate antigen-specific Tregs. Here, we compared CAR with endogenous T cell receptor (TCR)/CD28 activation in human Tregs. Strikingly, CAR Tregs displayed increased cytotoxicity and diminished suppression of antigen-presenting cells and effector T (Teff) cells compared with TCR/CD28 activated Tregs. RNA sequencing revealed that CAR Tregs activate Teff cell gene programs. Indeed, CAR Tregs secreted high levels of inflammatory cytokines, with a subset of FOXP3^+^ CAR Tregs uniquely acquiring CD40L surface expression and producing IFNγ. Interestingly, decreasing CAR antigen affinity reduced Teff cell gene expression and inflammatory cytokine production by CAR Tregs. Our findings showcase the impact of engineered receptor activation on Treg biology and support tailoring CAR constructs to Tregs for maximal therapeutic efficacy.

**Graphical Abstract:** 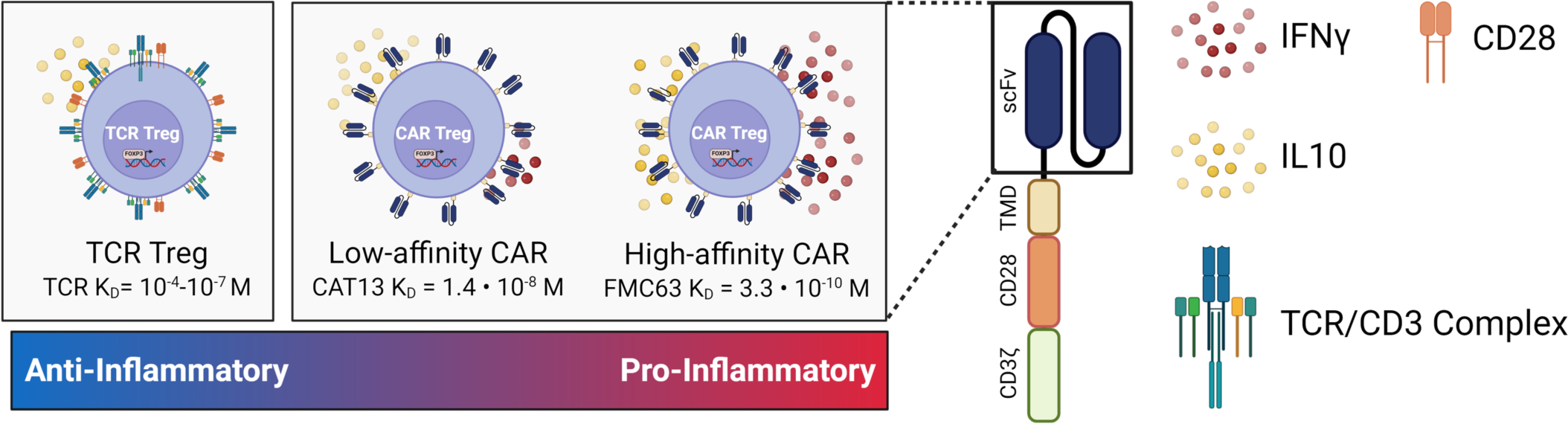

## INTRODUCTION

Recent advancements in transplantation medicine and autoimmune disorder treatments have generated optimism for more effective and long-lasting therapies. Nevertheless, a significant drawback persists in the dependency on broad immunosuppressive therapies that are accompanied by various systemic side effects and significantly burden patients, ranging from vulnerability to infections, cancer risk, hyperglycemia, and multi-organ damage to expensive lifelong treatments and severe long-term complications ^1–3^. As a result, the demand for localized antigen-specific immunomodulatory strategies has never been more urgent.

Regulatory T cells (Tregs), a small (3-6%) but indispensable subset of CD4^+^ T cells, have emerged as a potential cornerstone for such targeted interventions ^4,5^. Characterized by their unique cytokine and inhibitory receptor profiles and expression of the transcription factor FOXP3 ^6–8^, Tregs inhibit immune responses and promote tissue repair locally upon antigen recognition ^9,10^. However, Treg infusion in clinical settings for transplant and autoimmune disease has resulted in limited efficacy due to factors like antigen specificity, low abundance and expansion, functional instability upon *ex vivo* expansion, and limited *in vivo* survival ^5,11–13^.

The groundbreaking success of chimeric antigen receptor (CAR) technology in oncology has propelled interest in its application to Tregs. CARs are designer proteins comprising an extracellular antigen-binding domain, typically an antibody-derived single chain fragment variable (scFv), and an intracellular signaling domain, enabling T-cell activation by an antigen of choice ^14^. The success of CAR T cells in treating liquid tumors with unprecedented remission rates, with currently seven CAR T-cell therapies approved by the U.S. Food and Drug Administration (FDA) ^15^, has kindled interest in the generation of CAR Tregs to solve the problems of Treg antigen specificity and low numbers.

Initial results from CAR Tregs in preclinical humanized mouse models have shown promise in preventing graft-vs.-host disease and skin graft rejection ^16–19^. Yet, CAR Tregs have displayed lackluster efficacy as stand-alone agents in solid organ transplant rejection and autoimmune disease in immunocompetent murine and non-human primate models, as CAR Tregs required combination with immunosuppressive molecules to show efficacy ^20,21^ and were either ineffective or only shown to prevent, not reverse, autoimmune disease ^22–24^. In contrast, allo-antigen-specific murine Tregs suffice to prevent acute and chronic rejection of skin allografts in C57BL/6 mice ^25^ and murine T cell receptor (TCR) transgenic islet antigen-specific Tregs reverse autoimmune diabetes in non-obese diabetic (NOD) mice ^26^. Altogether, these preclinical data suggest that CAR Treg engineering and generation require further optimization for CAR Tregs to go from immunosuppressive drug adjuvants or partial replacements to an independent immunomodulatory intervention. Moreover, reports that CAR Tregs can be cytotoxic towards target cells ^27,28^ has also cast doubt on their safety and invites discussion on target selection for CAR Treg-mediated immune protection. Recently started and upcoming clinical trials testing CAR Tregs in organ transplantation add urgency to a preemptive investigation into CAR Treg therapy safety and limitations ^29,30^.

One plausible reason for the suboptimal performance of CAR Tregs lies in the fact that CAR constructs were originally designed and optimized for proinflammatory and cytotoxic T cells — a functional contradiction to the immunosuppressive nature of Tregs. T cell receptor (TCR) signaling is a complex cascade of events initiated by the engagement of the TCR with its cognate antigen-MHC complex on an antigen-presenting cell (APC), so called signal 1. Robust T-cell activation requires an additional input, costimulation, or signal 2, which is transmitted upon the binding of CD28 on the T-cell surface to CD80 or CD86 on the APC surface ^31^. Notably, the TCR itself does not participate in signal transduction, relying instead on the associated CD3 protein complex containing CD3δ, CD3ε, and CD3γ, each with one immunoreceptor tyrosine-based activating motif (ITAM) signaling domain, and CD3ζ, which contains three ITAMs and thus transduces the strongest signal ^32^. Strength and duration of this signaling ensemble orchestrate the functional outcomes of Treg activity, influencing their proliferation, immunosuppressive activity, and stability ^33–35^. TCR signaling operates via a network of kinases, adaptor molecules, and transcription factors, ensuring a highly regulated and specific immune response. Current CAR constructs attempt to mimic this by containing signal 1 (CD3) and signal 2 (CD28) within the CAR intracellular signaling domain, leading to their simultaneous activation upon engagement of the CAR scFv with its target antigen.

Previous literature has predominantly focused on the binary outcomes of CAR activation rather than delving into the nuanced functional outcomes of CAR Treg stimulation as compared to their TCR/CD28 stimulated counterparts. Such oversight could contribute to the observed suboptimal performance of CAR Tregs in preclinical settings, underlining the need for a comprehensive reevaluation. This study aims to bridge this gap, asking critical questions about the outcomes of CAR versus natural TCR/CD28 signaling in Tregs. Specifically, what intrinsic pathways might the current CAR constructs be missing or inappropriately triggering? By rigorously assessing these functional outcomes, we aim to optimize CAR Treg design, positioning it as a central element in the next generation of localized, antigen-specific immunomodulatory strategies.

Utilizing a variety of assays and techniques, we compared the activation, function, stability, and gene expression profiles of engineered CAR Tregs with those of naturally activated TCR/CD28 Tregs. Our investigation uncovered substantial alterations in Treg phenotype and function upon CAR-mediated activation, notably a shift towards a more inflammatory and cytotoxic gene expression profile and behavior. Indeed, we found *de novo* expression of CD40L as a surface marker associated with a subset of proinflammatory CAR Tregs. Finally, we identified scFv affinity as a CAR design parameter that modulates CAR Treg inflammatory cytokine production, with Treg activation via a lower affinity CAR resulting in a cytokine expression profile similar to that of TCR/CD28-activated Tregs.

## RESULTS

### Human CAR Treg generation

To systematically evaluate the phenotypic and functional discrepancies between chimeric antigen receptor (CAR) and endogenous T cell receptor (TCR)/CD28 mediated activation of human regulatory T cells (Tregs), we used a well-established anti-human CD19 CAR construct ^36^ with minor modifications, featuring an N-terminus Myc-tag to assess CAR surface expression, a CD28-CD3zeta signaling domain, and a green fluorescent protein (GFP) reporter gene to identify CAR-expressing cells (**Figure 1A**). We then magnetically isolated CD4^+^ T cells and CD8^+^ T cells from human peripheral blood (**Figure 1B**) and used fluorescence assisted cell sorting (FACS) to further purify CD4^+^CD25^hi^CD127^low^ Tregs ^37,38^ and CD4^+^CD25^low^CD127^hi^ effector T (Teff) cells from the CD4^+^ T cells (**Figure 1C**). Isolated cells were activated with anti-CD3/CD28 beads and interleukin 2 (IL-2), transduced with CAR lentivirus two days later, and expanded in the presence of IL-2. As expected, Tregs co-expressed the Treg lineage transcription factors FOXP3 and HELIOS ^12,16^, whereas Teff cells did not (**Figure 1C**). CAR-expressing cells were isolated by FACS based on GFP expression (**Figure 1D**) and CAR surface expression on the isolated cells confirmed using flow cytometry (**Figure 1D**). Expanded CAR Tregs were used for experiments 9-12 days after cell isolation from peripheral blood (**Figure 1B**).

**Figure 1.**
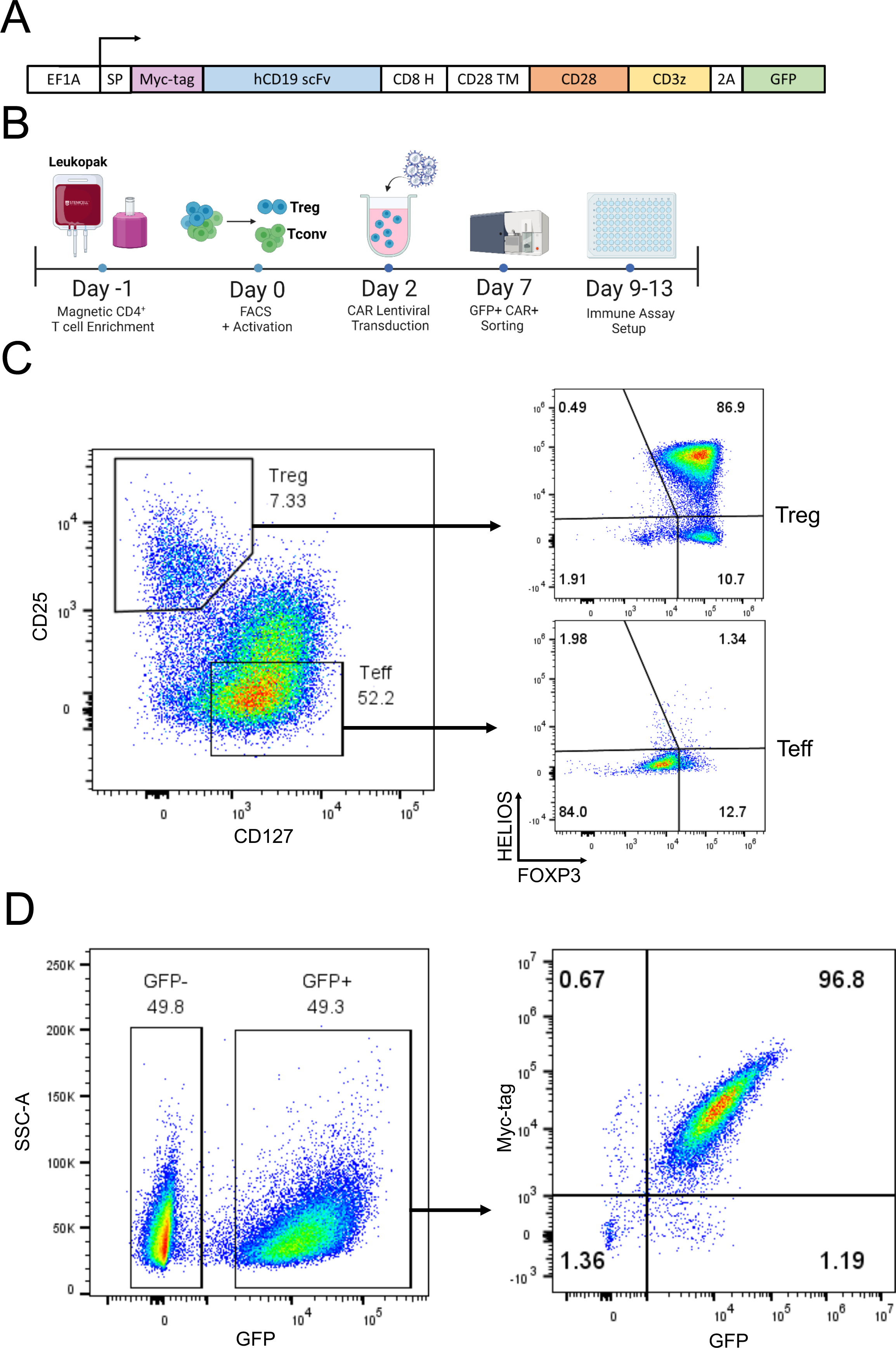
Human CAR Treg generation. (A) Schematic of chimeric antigen receptor (CAR) constructs used in this study. (B) Workflow to isolate human CD4^+^ regulatory T cells (Tregs) and effector T cells (Teff), introduce a CAR, expand, and sort CAR-expressing cells for immune assays. (C) Representative dot plots of Treg sorting strategy with CD25^hi^CD127^low^ Tregs and CD25^low^CD127^hi^ Teff on the left and Treg phenotype assessment with FOXP3^+^HELIOS^+^ Tregs and FOXP3^−^HELIOS^−^ Teff cells on the right. (D) Representative dot plots of Treg transduction efficiency with CD19CAR-2A-GFP lentivirus, based on GFP expression on the left and CAR surface expression (Myc-tag) and reporter gene expression (GFP) after sorting GFP^+^ cells on the right.

### CAR Tregs are functionally distinct from TCR/CD28 activated Tregs

To accurately model endogenous TCR immune synapses, endogenous CD28 engagement, and CAR immune synapses and to reduce confounding factors when comparing CAR and TCR/CD28 activation, we generated target cell lines to elicit TCR/CD28 and CAR activation. Specifically, we transduced either a CD64-2A-CD80 or a CD19 extracellular domain fused to a platelet-derived growth factor (PDGFR) transmembrane transgene into K562 cells, a human myelogenous leukemia cell line that lacks HLA, CD80, and CD86 expression and thus does not activate T cells. CD64 is a high-affinity Fc receptor and CD80 binds to CD28. CD64-expressing K562 cells were loaded with anti-CD3 antibody, as previously described, to activate Tregs via the TCR ^39^. Expanded CAR Tregs were incubated with irradiated K562 cells (no activation, “No Act”), CD64-CD80-K562 cells tagged with anti-CD3 antibody (TCR/CD28 activation) or CD19-K562 cells (CAR activation) (**Figure 2A**).

**Figure 2.**
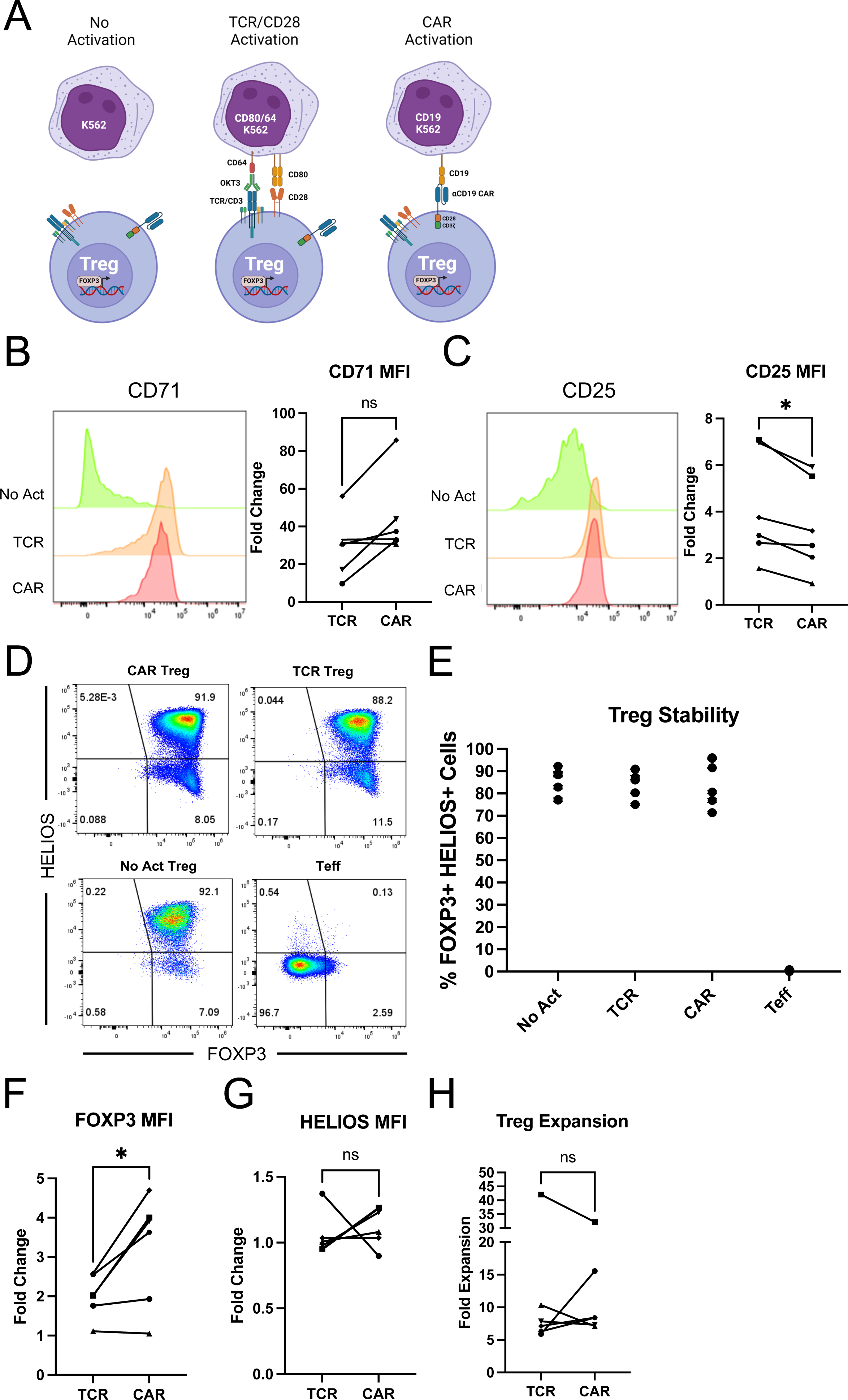
CAR and TCR/CD28 activation result in phenotypically similar Tregs. (A) Schematic with the three modes of activation used in this study: No Activation with target K562 cells (No Act), TCR/CD28 activation with target K562 cells expressing CD64 loaded with anti-CD3 antibody and CD80 (TCR), and CAR activation with target K562 cells expressing CD19 (CAR). (B) CD71 surface expression 48h after Treg activation. Representative histogram on the left and summary data across donors of fold change in CD71 mean fluorescence intensity (MFI) in relation to No Act Tregs on the right. (C) CD25 surface expression 48h after Treg activation. Representative histogram on the left and summary data across donors of fold change in CD25 MFI in relation to No Act Tregs on the right. (D) Representative dot plots of FOXP3 and HELIOS expression in CAR Treg, TCR Treg, and No Act Treg, as well as in Teff cells as a negative staining control. (E) Percentage of FOXP3^+^HELIOS^+^ cells across activation modes and donors. (F) Fold change in FOXP3 MFI in TCR Tregs or CAR Tregs over No Act Tregs across donors. (G) Fold change in HELIOS MFI in TCR Tregs or CAR Tregs over No Act Tregs across donors. (H) Fold expansion in cell number for TCR Tregs and CAR Tregs one-week post-activation. For Figures 2B, 2C, 2F, 2G, and 2H, values represent mean ± SD of technical triplicates per blood donor, with lines collecting the data points from the same donor. Unpaired Student’s t test. *, p < 0.05; ns, not significant.

Our first aim was to investigate whether stimulation via a CAR or endogenous TCR/CD28 pathways results in different levels of Treg activation. Given the higher affinity of CARs, including the FMC63 scFv-based CD19 CAR used here ^36,40^, compared to TCRs ^41,42^, we hypothesized that CAR-mediated activation would lead to a heightened activation state. To assess this, CAR Tregs were coincubated with each K562 cell line, harvested after 48 hours, and their activation status was evaluated by measuring the cell surface expression of CD71 (transferrin receptor), a well-established early marker of T-cell activation. Interestingly, no statistically significant difference was found in the mean fluorescence intensity (MFI) of CD71 between CAR- and TCR/CD28-activated Tregs across blood donors (**Figure 2B**).

In parallel, we examined the expression of CD25, the high affinity alpha chain of the IL-2 receptor. In addition to being a T-cell activation marker, CD25 is constitutively expressed by Tregs and is crucial for their immunosuppressive function via IL-2 sequestration ^4,43^. We were intrigued to find that TCR/CD28-activated Tregs had slightly but significantly higher levels of CD25 expression compared to CAR-activated Tregs after 48 hours of coculture (**Figure 2C**).

Next, we assessed the stability of the Treg phenotype on day 8 post-activation, as Tregs can convert into effector-like cells under certain conditions, such as highly inflammatory microenvironments and repeated *in vitro* stimulation ^44–46^. To gauge this, we assessed the expression of the Treg lineage transcription factors FOXP3 and HELIOS. FOXP3 is indispensable for Treg identity and function ^6–8^, while HELIOS is believed to confer stability to the Treg phenotype ^47^. Across blood donors, we found that all activation conditions maintained a distinct (**Figure 2D**) and equally abundant (**Figure 2E**) FOXP3^+^HELIOS^+^ cell population, indicating that neither CAR nor TCR/CD28 activation led to Treg destabilization. Nevertheless, FOXP3 levels were higher in CAR versus TCR/CD28 activated Tregs (**Figure 2F**), whereas HELIOS levels were similar (**Figure 2G**).

To complete this initial phenotypic characterization, we evaluated the cells’ expansion capacity – a critical attribute considering the current challenges in achieving therapeutically sufficient Treg numbers for infusion ^12^. In line with activation and stability, expansion of CAR and TCR/CD28-activated Tregs was similar across donors (**Figure 2H**).

While phenotypic characterization indicated that CAR-activated Tregs closely resemble TCR/CD28-activated Tregs, functional assays are essential to characterize these modes of activation. Tregs have an arsenal of over a dozen known suppressive mechanisms, inhibiting immune responses both through contact-independent pathways – such as the sequestration of IL-2 via CD25 and the secretion of anti-inflammatory cytokines such as IL-10 – and contact-dependent pathways, such as CTLA4-mediated trogocytosis of costimulatory molecules CD80 and CD86 from APCs ^4,9^.

To delineate how CAR activation influences these functionalities compared to endogenous TCR/CD28 activation, we first employed a modified *in vitro* T-cell suppression assay where Tregs were activated via CAR or TCR/CD28 overnight and then co-incubated with CellTrace dye-labeled CD4^+^ and CD8^+^ T responder (Tresp) cells activated with anti-CD3/CD28 beads overnight in parallel at different Treg to Tresp cell ratios ^48,49^. Interestingly, CAR-activated Tregs were less efficacious than their TCR/CD28-activated counterparts in inhibiting CD4^+^ (**Figure 3A**) and CD8^+^ (**Figure 3B**) Tresp cell proliferation. Additionally, to assess APC modulatory activity, we co-incubated Tregs with NALM6, a CD19^+^ B-cell leukemia cell line; CAR Tregs were incubated with NALM6 and untransduced Tregs with CD80-CD64-NALM6 loaded with anti-CD3 antibody to test CAR activation and TCR/CD28 activation, respectively. Four days later, CD80 surface expression was measured by flow cytometry ^50^. Consistent with our observations on T-cell suppression (**Figures 3A and 3B**), CD80 expression on the target cells was downregulated to a lesser extent by CAR Tregs than by their TCR/CD28-activated counterparts (**Figure 3C**). However, the same trend was not observed when using primary CD14^+^ monocyte-derived dendritic cells (moDCs) as target APCs. Irrespective of the form of activation, all Treg conditions downregulated CD80 (**Figure S1A**) and CD86 (**Figure S1B**) on moDCs to the same extent.

**Figure 3.**
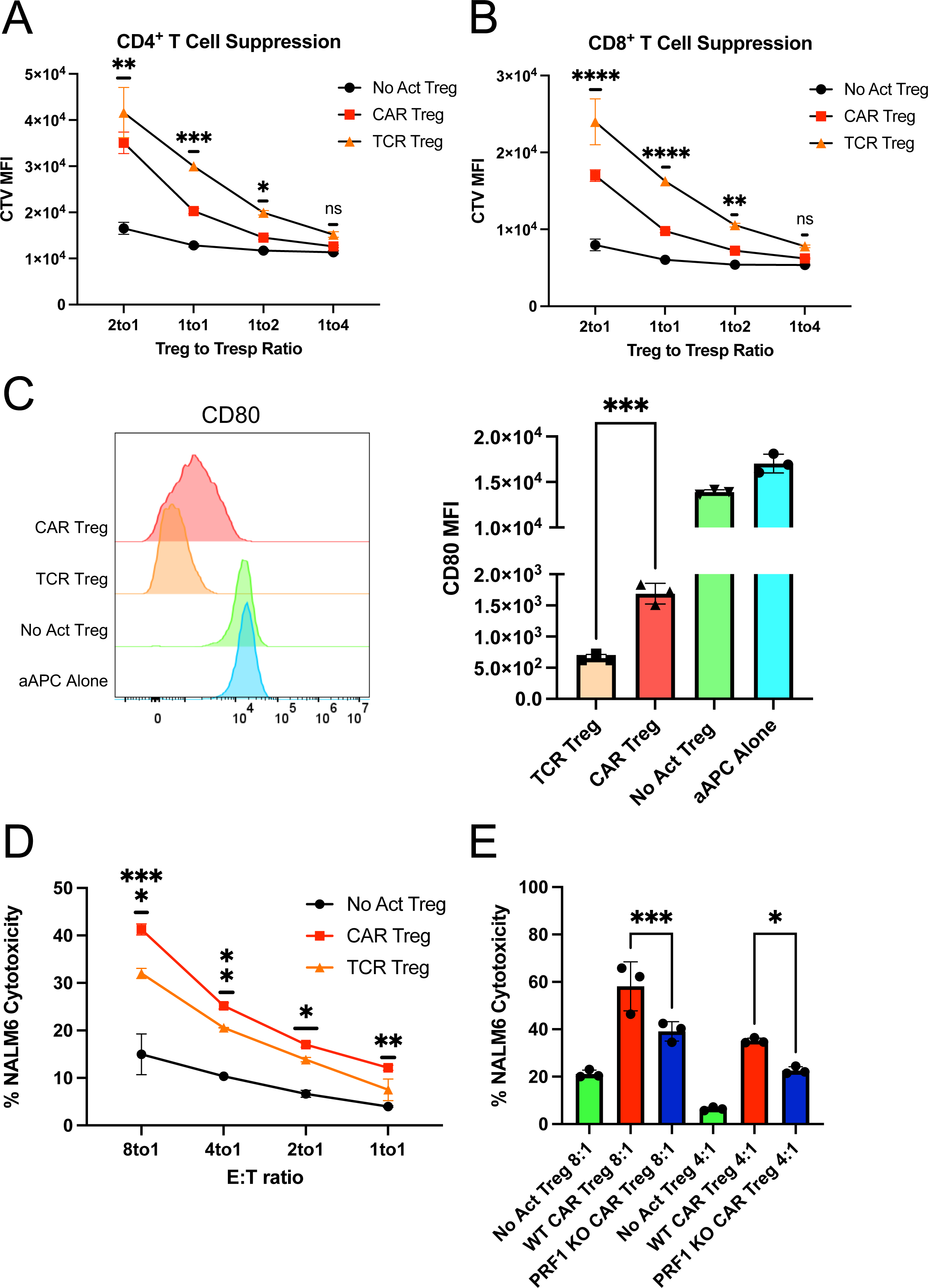
CAR activation leads to a shift from suppression to cytotoxicity in Tregs. (A) Inhibition of CellTrace Violet (CTV) labeled CD4^+^ T responder cell (Tresp) proliferation by Tregs. (B) Inhibition of CTV labeled CD8^+^ Tresp proliferation by Tregs. (C) Downregulation of CD80 surface expression in CD80-NALM6 cells (aAPC – artificial antigen presenting cells) by Tregs. Representative histograms on the left and summary data on the right. (D) Treg cytotoxicity towards target NALM6 cells at different effector to target (E:T) ratios. (E) WT and PRF1 KO CAR Treg cytotoxicity towards target NALM6 cells at different E:T ratios. Values represent technical replicates of representative experiments. Bars represent mean ± SD. One-way ANOVA test with Tukey’s multiple comparison correction. ****, p < 0.0001; ***, p < 0.001; **, p < 0.01; *, p < 0.05; ns, not significant.

Despite not being as studied as other Treg suppressive strategies, Tregs have been found to suppress immune responses via direct cytotoxicity. The most common mechanism of cytotoxicity by T cells and NK cells is the perforin/granzyme pathway, where perforin forms pores in the membrane of the target cells, allowing the delivery of granzymes into the target cells and subsequent induced cell death ^51^. Tregs have been shown to kill their target cells via the perforin/granzyme pathway, with both granzyme B and perforin being required for optimal Treg-mediated suppression by either eliminating APCs or CD8^+^ T cells and natural killer (NK) cells directly ^52–55^. Considering that CAR signaling was initially designed for triggering inflammatory responses and cytotoxicity by Teff cells, we hypothesized that CAR Tregs might be more cytotoxic than TCR/CD28-activated Tregs. To test this, we again incubated CAR Tregs with NALM6 and untransduced Tregs with CD80-CD64-NALM6 loaded with anti-CD3 antibody to test CAR activation and TCR/CD28 activation, respectively. In agreement with our hypothesis, CAR Tregs were significantly more cytotoxic than TCR/CD28-activated Tregs towards NALM6 cells at different effector to target (E:T) ratios (**Figure 3D**). In contrast, CAR Teff and TCR/CD28-activated Teff cells killed NALM6 cells to a similar extent (**Figure S1C**). To investigate whether CAR Treg cytotoxicity depends on the perforin/granzyme pathway, we deleted the PRF1 gene, which encodes perforin, in CAR Tregs using CRISPR/Cas9 and tested the cytotoxicity of the resulting cells towards NALM6 cells. Indeed, PRF1 knockout (KO) CAR Tregs (59% indel efficiency by Tracking of Indels by Decomposition – TIDE – analysis ^56^) were less effective at killing NALM6 cells than their WT counterparts (**Figure 3E**). Additionally, we investigated whether CAR Tregs could eliminate non-immune cells. Most CAR Treg therapies being currently investigated directly target the tissues to be protected from immune rejection ^29^ and hence it is fundamental to ask whether CAR Tregs protect the targeted tissue rather than participating in its elimination. To answer this question, we ectopically expressed our CD19 extracellular domain fused to a PDGFR transmembrane transgene in A549 lung cancer epithelial cells. Interestingly, CAR Tregs were not cytotoxic towards CD19-A549 cells, in contrast with CAR Teff cells (**Figure S1D**).

### CAR activation alters the natural transcriptome of Tregs

Given our observations on CAR-activated Tregs’ enhanced cytotoxicity and reduced suppressive function in comparison with TCR/CD28-activated Tregs, a crucial question emerged: Why do these alterations occur? Answering this question holds significance not only for our understanding of Treg biology but also for the efficacy of CAR Tregs in the clinic. To address this question, we co-incubated CAR Tregs and CAR Teff cells with each of the three types of target K562 cell lines for no activation (“No Act”), TCR/CD28 activation (“TCR”), and CAR activation (“CAR”) and performed RNA sequencing on CD4^+^ T cells isolated 24 hours post-activation. Whole-transcriptome analysis with two blood donors under all six conditions revealed that both CAR and TCR/CD28 activated Tregs upregulated NR4A1 and NR4A3, which are immediate-early genes induced by TCR signaling ^57^; IL10 and EBI3, which encode the anti-inflammatory cytokines IL-10 and IL-35 ^58,59^, respectively; CCR8, a chemokine receptor gene expressed in highly activated Tregs ^60^; and IL1R2, a gene that encodes a decoy receptor for the inflammatory cytokine IL-1 ^61^ **(Tables S1 and S2)**. However, 3,680 genes were upregulated by CAR activation in Tregs, while only 1,236 genes were upregulated in response to TCR/CD28 activation, suggesting that CAR activation elicits a more pronounced transcriptional response in Tregs than does physiological TCR/CD28 signaling. Of note, a similar pattern was observed in Teff cells, with CAR activation upregulating 4,013 genes compared to 2,058 genes with TCR/CD28 activation **(Tables S3 and S4)**. In addition, CAR Treg and CAR Teff cells clustered closest together despite being different cell types (**Figure 4A**). In line with this observation, joint analysis of genes upregulated by CAR Tregs, TCR Tregs, CAR Teff, and TCR Teff in comparison with the respective non-activated cells revealed that CAR Tregs shared 1,038 upregulated genes uniquely with CAR Teff but only 219 upregulated genes uniquely with TCR Tregs (**Figure 4B**). These findings suggested that CAR activation induces the expression of Teff cell gene programs in Tregs, as if CAR signaling partly overrides intrinsic Treg gene programs. Indeed, the top differentially expressed protein-coding genes between CAR Tregs and TCR Tregs (**Table S5**) included key proinflammatory cytokine and chemokine genes, such as IFNG, IL17F, IL3, CCL3, CCL19, and CSF3 (**Figure 4C**). Gene Set Enrichment Analysis (GSEA) ^62^ revealed that the upregulated gene programs in CAR Tregs in comparison to those in TCR Tregs were primarily those related to cytokine signaling and inflammation, such as PI3K-AKT signaling, IL-17 signaling, cytokines and inflammatory response, and proinflammatory and profibrotic mediators (**Figure 4D, Figure S2A**), with CAR Tregs expressing higher levels of proinflammatory cytokine and chemokine genes than TCR Tregs (**Figure S2B**). Curiously, CAR activation also resulted in differences in chemokine receptor gene expression: while the expression of CCR2 and CCR5, high in TCR Tregs, was even lower in CAR Tregs than in CAR Teff and TCR Teff cells, CCR8 expression, absent in Teff cells, remained as high in CAR Tregs as in TCR Tregs (**Figure S3**).

**Figure 4.**
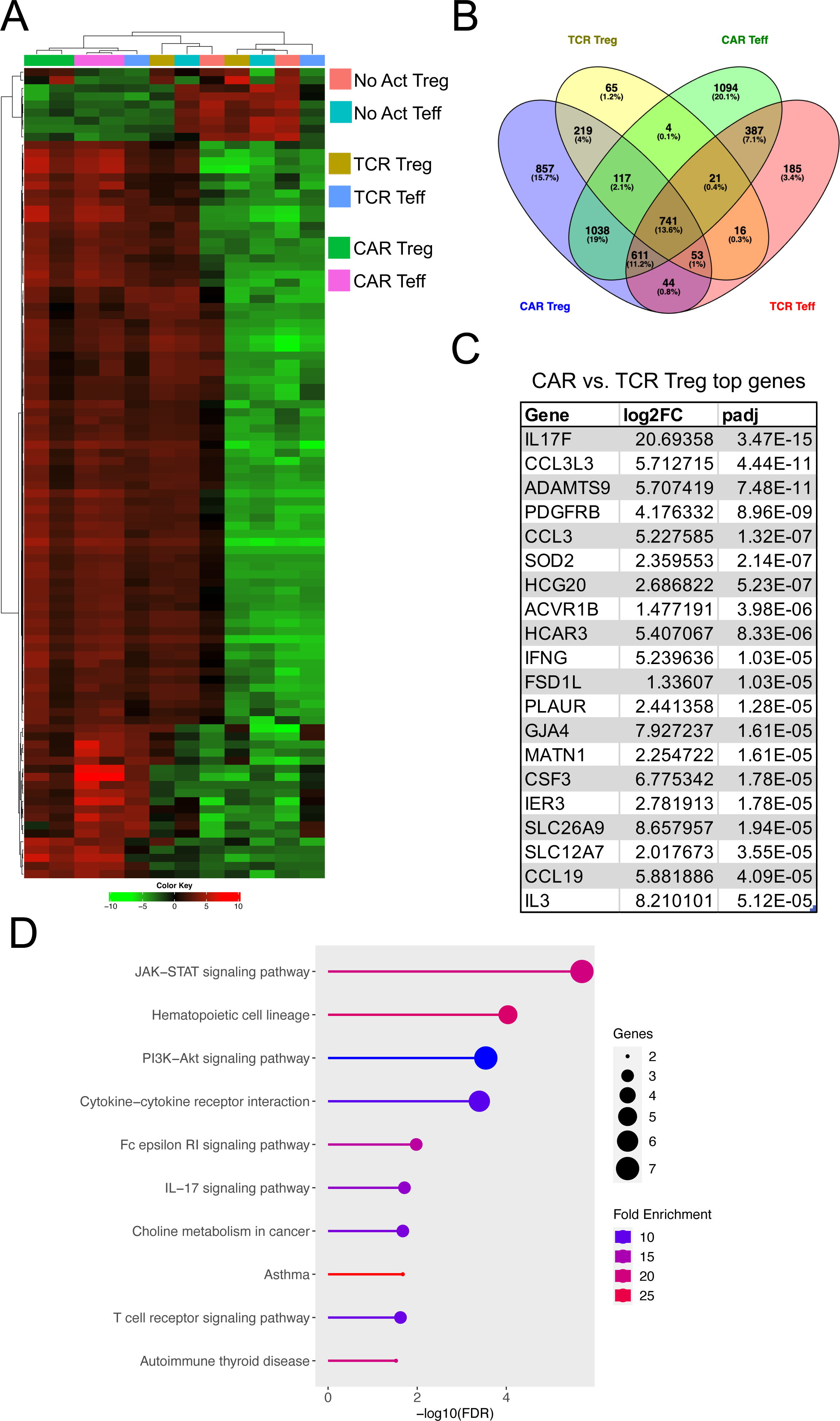
CAR activation induces pro-inflammatory gene programs in Tregs. (A) Heatmap clustered by column (sample) and by row (gene) with top 100 most differentially expressed genes between No Act Tregs, TCR Tregs, CAR Tregs, No Act Teff, TCR Teff, and CAR Teff. (B) Venn diagram with genes upregulated in TCR Tregs, CAR Tregs, TCR Teff, and CAR Teff in relation to their respective No Act cell types. Number of genes and respective percentage of the total number of genes are indicated in each intersection. (C) Top 20 protein-coding genes most differentially expressed in CAR Tregs compared with TCR Tregs. FC, fold change; padj, adjusted p value. (D) KEGG pathway gene set enrichment analysis (GSEA) of CAR Tregs vs. TCR Tregs. FDR, false discovery rate.

### CAR activation induces a distinct cytokine production pattern in Tregs

Considering the marked increased in pro-inflammatory cytokine and chemokine gene expression by CAR Tregs compared to TCR/CD28-activated Tregs, we sought to validate this pattern at the protein level. First, we collected the supernatants of 48h co-cultures of CAR Tregs and CAR Teff cells with irradiated K562 cells (no activation), CD64-CD80-K562 cells with anti-CD3 (TCR/CD28 activation) or CD19-K562 cells (CAR activation) for cytokine quantitation using multiplex enzyme-linked immunosorbent assay (ELISA). CAR Tregs secreted more shed CD40L (sCD40L), IFNγ and IL-17A, while secreting same amount of TNFα and IL-10 and more IL-13 than TCR Tregs (**Figure 5A**). CAR Tregs also secreted more IL-3, G-CSF, IL-4, IL-6, and TNFβ than TCR Tregs (**Figure S4**). Overall, these cytokine secretion data echoed our RNA-seq data, suggesting that CAR activation leads to notably higher inflammatory cytokine and chemokine production in Tregs while maintaining immunosuppressive cytokine secretion levels. One of the most intriguing findings from the cytokine quantification was IFNγ secretion by CAR Tregs, reaching levels comparable to those of CAR Teff and TCR Teff cells (**Figure 5A**), in line with IFNG being one of the most differentially expressed genes between CAR Tregs and TCR Tregs (**Figure 4C**). Even though our Treg lineage stability analysis indicated that CAR-activated Tregs retained FOXP3 and HELIOS expression to the same extent as TCR/CD28-activated Tregs (**Figures 2D and 2E**), we set out to examine whether the high IFNγ levels measured using bulk RNA-seq and ELISA were the product of contaminating Teff cells and/or FOXP3 negative ex-Treg cells. We performed intracellular cytokine staining for CAR Tregs and CAR Teff cells following no activation, CAR activation, or TCR/CD28 activation with the respective target K562 cell lines overnight followed by 5 hours of brefeldin A and found that CAR-activated FOXP3^+^ Tregs, but not TCR/CD28-activated or resting Tregs, produced IFNγ (**Figure 5B**), suggesting that CAR Tregs do not become unstable and lose Treg identity prior to producing IFNγ. In line with this hypothesis, Tregs did not produce IL-2 regardless of activation mode (**Figure 5C**), a key hallmark of Treg identity ^63^. In contrast, Teff cells produced IFNγ (**Figure 5B**) and IL-2 (**Figure 5C**) when activated via CAR or endogenous TCR/CD28, as expected. Therefore, CAR activation generates a unique subset of Tregs that are proinflammatory yet retain key Treg identity markers. This implies that CAR activation is leading to the emergence of a functionally distinct Treg subpopulation that can potentially influence the balance of immune responses in novel ways.

**Figure 5.**
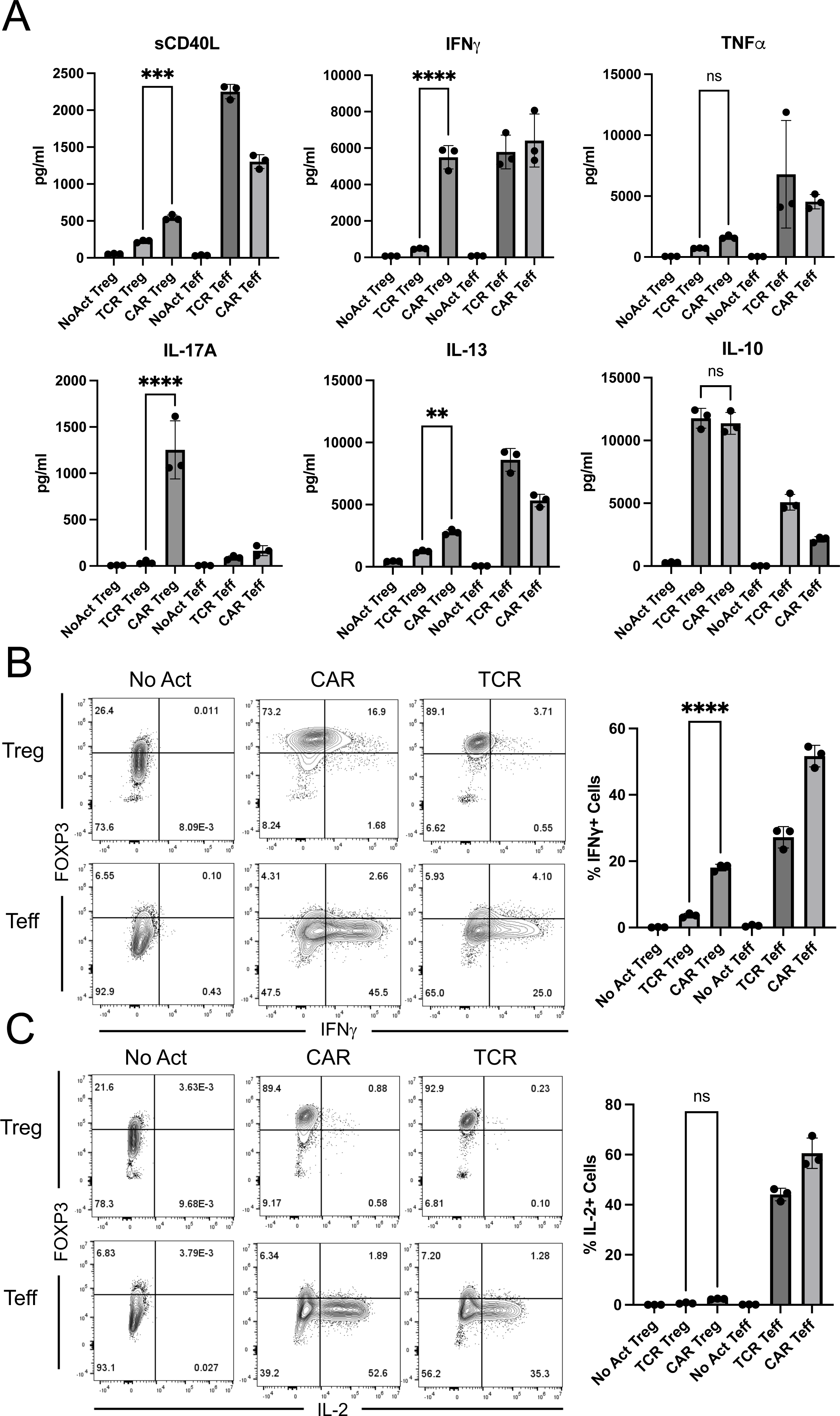
CAR Tregs uniquely produce inflammatory cytokines. (A) Levels of cytokines secreted into the supernatant by No Act Tregs, TCR Tregs, CAR Tregs, No Act Teff, TCR Teff, and CAR Teff 48h post-activation. (B) Intracellular levels of IFNG produced by No Act Tregs, TCR Tregs, CAR Tregs, No Act Teff, TCR Teff, and CAR Teff 18h post-activation. Representative countour plots on the left and summary data on the right. (C) Intracellular levels of IL-2 produced by No Act Tregs, TCR Tregs, CAR Tregs, No Act Teff, TCR Teff, and CAR Teff 18h post-activation. Representative contour plots on the left and summary data on the right. Values represent technical replicates of representative experiments. Bars represent mean ± SD. One-way ANOVA test with Tukey’s multiple comparison correction. ****, p < 0.0001; ***, p < 0.001; **, p < 0.01; *, p < 0.05; ns, not significant.

### Characterizing the proinflammatory CAR Treg subset

As we delved deeper into understanding CAR Tregs’ unique functional attributes, we recognized the importance of investigating cell surface markers. In addition to being important phenotypic signposts, surface markers can be used to better identify and purify cell subsets, allowing for a more nuanced understanding of CAR Tregs. Upon scrutinizing our RNA-seq data, specifically the genes upregulated in different modes of activation (CAR vs. TCR/CD28) and cell types (Treg vs. Teff) (**Figure 4B**), we noticed that CAR Tregs, CAR Teff, and TCR Teff, but not TCR Tregs, upregulated CD40LG (**Table S6**), a gene coding for the well-known Teff cell activation marker CD40L or CD154 ^64^. In addition, CAR Tregs secreted significantly more sCD40L than TCR/CD28-activated Tregs (**Figure 5A**). Conversely, TCR Tregs, but not any of the other 3 activated conditions, upregulated FCRL3 and ENTPD1 (**Table S7**). ENTPD1 encodes CD39, a cell surface ectoenzyme expressed in Tregs that converts ATP into the immunosuppressive molecule adenosine^65^. Yet, TCR Tregs also uniquely upregulated ENTPD1-AS1 (**Table S7**), an anti-sense RNA previously shown to decrease CD39 expression ^66^. FCRL3, on the other hand, has been associated with TIGIT and HELIOS expression in Tregs ^67^. TIGIT is a surface marker expressed by Tregs that are highly suppressive towards Th1 cells, which secrete IFNγ, and Th17 cells, which secrete IL-17 ^68^. Molecularly, TIGIT is thought to induce phosphatase activity to downmodulate TCR signaling in the TIGIT-expressing Treg and to induce IL-10 production by dendritic cells upon binding to PVR on the surface of the dendritic cell ^69^. Although not statistically significant (p > 0.05), TCR/CD28-activated Tregs upregulated TIGIT transcript (**Table S1**), whereas CAR-activated Tregs did not (**Table S2**).

We then aimed to validate whether CD40L and TIGIT were differentially expressed in CAR- and TCR/CD28-activated Tregs at the surface protein level using flow cytometry, possibly offering a further detailed characterization of the unique pro-inflammatory CAR Treg phenotype. Following 48-hour activation, CAR Tregs displayed significantly higher CD40L and reduced TIGIT levels compared with TCR Tregs (**Figure 6A**), trends that were maintained one week after activation (**Figure 6B**). A targeted gene expression survey using quantitative polymerase chain reaction (qPCR) following 24-hour activation confirmed that CAR Tregs express higher levels of the Teff cell genes IFNG, GZMB, and CD40LG, and lower levels of TIGIT than TCR/CD28-activated Tregs (**Figure 6C**). Yet, CAR Tregs did not express higher levels of TBX21, GATA3, or RORC, genes coding for the master transcription factors of the main CD4^+^ Teff cell lineages Th1, Th2, and Th17, respectively, or STAT1, a key transcription factor in IFNγ signaling ^70,71^ (**Figure 6C**).

**Figure 6.**
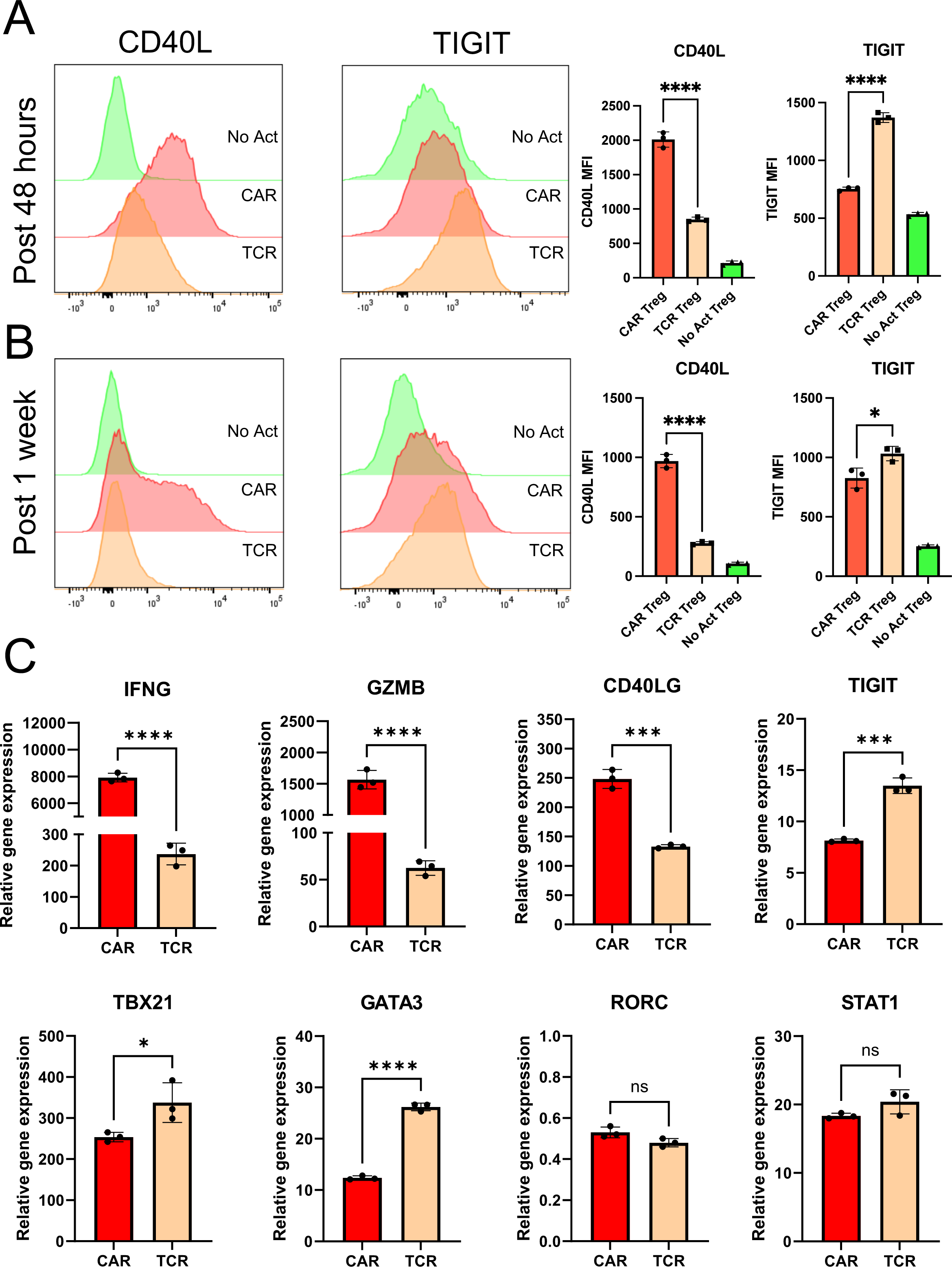
CAR activation induces CD40L expression in Tregs. (A) CD40L and TIGIT surface expression on No Act Tregs, TCR Tregs, and CAR Tregs 48h post-activation. Representative histograms on the left and summary data on the right. (B) CD40L and TIGIT surface expression on No Act Tregs, TCR Tregs, and CAR Tregs one-week post-activation. Representative histograms on the left and summary data on the right. (C) Expression of selected genes in CAR Tregs and TCR Tregs 24h post-activation, evaluated by qPCR. Values represent technical replicates of representative experiments. Bars represent mean ± SD. For Figures 6A and 6B, one-way ANOVA test with Tukey’s multiple comparison correction. For Figure 6C, unpaired Student’s t test. ****, p < 0.0001; ***, p < 0.001; **, p < 0.01; *, p < 0.05; ns, not significant.

### Lowering CAR affinity reduces inflammatory cytokine production by CAR Tregs

T-cell activation and function are influenced by the affinity of the TCR and the strength of costimulation ^72,73^. Moreover, as previously mentioned, Tregs exhibit dampened activation of several pathways downstream of TCR signaling ^34,35^. Inspired by these notions, we modified our CAR construct to dissect which of its features was responsible for the proinflammatory shift observed in CAR-activated Tregs and potentially better mimic endogenous TCR/CD28 engagement in Tregs. To reduce affinity, we modified the extracellular domain of the CAR by swapping the FMC63 scFv domain with an scFv sequence, CAT-13.1E10, which binds to the same CD19 residues as FMC63 but with a 40-fold lower affinity ^40^. To reduce costimulation strength, we modified the intracellular domain of the CAR by mutating all tyrosines of the CD28 signaling domain, as well as both prolines of its PYAP domain, which binds to Lck ^31,74^. We then introduced these two new CARs, which we called CAT and PY3, respectively, into Tregs to investigate the impact of affinity and costimulation strength on CAR Tregs. We activated CAR, CAT, and PY3 Tregs via the CAR with irradiated CD19-K562 cells (in parallel with TCR/CD28 activation and no activation) and performed whole-transcriptome RNA-seq as described earlier. We found that CAR, CAT, and PY3 Tregs clustered together and TCR and No Act Tregs clustered together based on gene expression (**Figure S5A**), indicating that, at the whole transcriptome level, activation via a lower affinity CAR or a lower signal 2 strength CAR remain more akin to CAR activation than to endogenous TCR/CD28 activation. Nevertheless, looking at the genes uniquely upregulated by each of these four modes of activation (TCR, CAR, CAT, PY3) revealed that CAR Tregs upregulated more genes uniquely (1,394) than any of the other conditions (**Figure S5B**). Focusing on CAT Tregs and PY3 Tregs, we found that, despite a large overlap in upregulated genes between these two conditions (**Figure S5C**), PY3 Tregs uniquely upregulated the inflammatory genes IL17A, IL1B, CXCL11, CSF3, and, importantly, CD40LG, as well as the cytotoxicity genes GZMB, CRTAM, and NKG7 (**Table S8**). Indeed, PY3 Tregs had IL17A, IFNG, CD40LG, and GZMB expression levels almost as high as CAR Tregs, whereas CAT Tregs had expression levels of these same genes almost as low as TCR/CD28-activated Tregs (**Figure S5D**). Interestingly, however, both CAT and PY3 Tregs still had CCR2, CCR5, and CXCR3 expression levels as low as CAR Tregs, suggesting that lower affinity (CAT) and lower costimulation strength (PY3) did not rescue expression of these chemokine receptor genes to the levels observed in TCR/CD28-activated Tregs (**Figure S5D**). Altogether, activation via the lower affinity CAT construct, but not via the lower costimulation strength PY3, resulted in visibly lower expression of inflammatory genes, kindling our interest in further comparing the CAR and CAT constructs head-to-head (**Figure 7A**). CAR and CAT Tregs had equivalent receptor surface expression post GFP^+^ cell sorting, based on Myc-tag expression (**Figure 7B**), and expanded to a similar extent post activation with irradiated CD19-K562 cells (**Figure 7C**). Yet, CAT Tregs upregulated CD71 to a smaller extent than CAR Tregs (**Figure 7D**). Importantly, activated CAR Tregs and CAT Tregs had an equally stable Treg phenotype, based on similar levels of CD25 (**Figure 7E**), FOXP3, and HELIOS (**Figures 7F-I**) expression. At the functional level, CAT Tregs were superior at suppressing CD4^+^ T cells (**Figure 8A**), but not CD8^+^ T cells (**Figure 8B**), downregulated CD80 surface expression on target cells to a larger extent (**Figure 8C**), and were less cytotoxic towards NALM6 cells (**Figure 8D**) than CAR Tregs. Moreover, CAT Tregs secreted sCD40L, IFNγ, TNFα, and IL-17A (**Figure 9**), as well as IL-3, IL-4, and IL-6 (**Figure S6**) at the same low levels as TCR/CD28-activated Tregs. Altogether, reducing the affinity of the CAR construct by 40-fold resulted in engineered Tregs with higher suppressive capacity, lower cytotoxic activity, and reduced inflammatory cytokine secretion.

**Figure 7.**
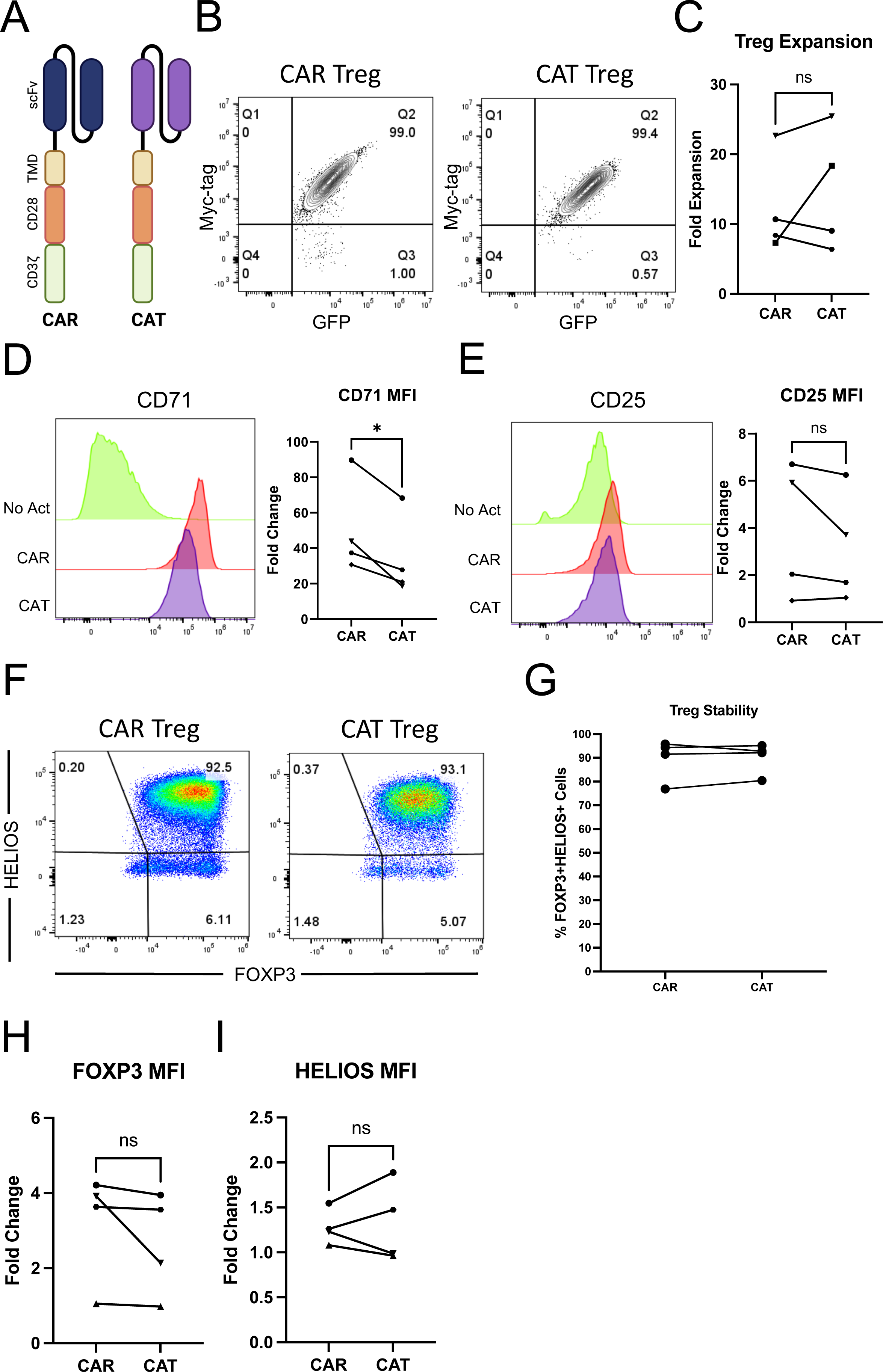
Lowering CAR affinity reduces the extent of CAR Treg activation. (A) Schematic of high affinity FMC63 CD19 CAR (CAR) and low affinity CAT-13.1E10 CD19 CAR (CAT). (B) Representative contour plot of surface expression (Myc-tag) of CAR and CAT constructs on Tregs. (C) Fold expansion in cell number for CAR Tregs and CAT Tregs one-week post-activation. (D) CD71 surface expression 48h after Treg activation. Representative histogram on the left and summary data across donors of fold change in CD71 mean fluorescence intensity (MFI) in relation to No Act Tregs on the right. (E) CD25 surface expression 48h after Treg activation. Representative histogram on the left and summary data across donors of fold change in CD25 MFI in relation to No Act Tregs on the right. (F) Representative dot plots of FOXP3 and HELIOS expression in CAR Tregs and CAT Tregs. (G) Percentage of FOXP3^+^HELIOS^+^ in CAR Tregs and CAT Tregs across donors. (H) Fold change in FOXP3 MFI in CAR Tregs and TCR Tregs over No Act Tregs across donors. (I) Fold change in HELIOS MFI in CAR Tregs and CAT Tregs over No Act Tregs across donors. For Figures 3C, 3D, 3E, 3G, 3H, and 3I, values are the mean ± SD of technical triplicates per blood donor, with lines collecting the data points from the same donor. Unpaired Student’s t test. *, p < 0.05; ns, not significant.

**Figure 8.**
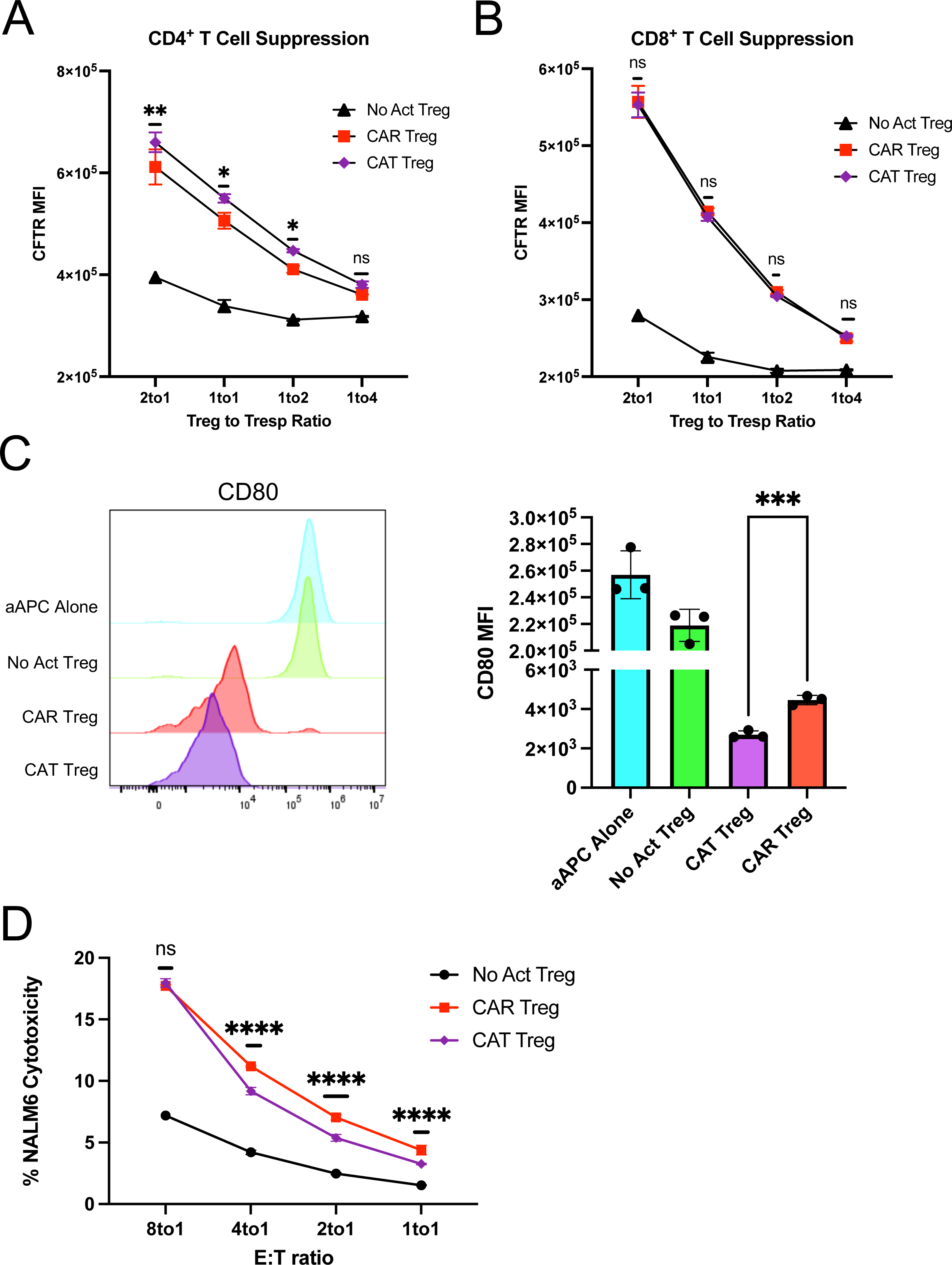
Lowering CAR affinity improves CAR Treg suppressive function. (A) Inhibition of CellTrace Far Red (CTFR) labeled CD4^+^ T responder cell (Tresp) proliferation by Tregs. (B) Inhibition of CTFR labeled CD8^+^ Tresp proliferation by Tregs. (C) Downregulation of CD80 surface expression in CD80-NALM6 cells (aAPC – artificial antigen presenting cells) by Tregs. Representative histograms on the left and summary data on the right. (D) Treg cytotoxicity towards target NALM6 cells at different effector to target (E:T) ratios. (E) WT and PRF1 KO CAR Treg cytotoxicity towards target NALM6 cells at different E:T ratios. Values represent technical replicates of representative experiments. Bars represent mean ± SD. One-way ANOVA test with Tukey’s multiple comparison correction. ****, p < 0.0001; ***, p < 0.001; **, p < 0.01; *, p < 0.05; ns, not significant.

**Figure 9.**
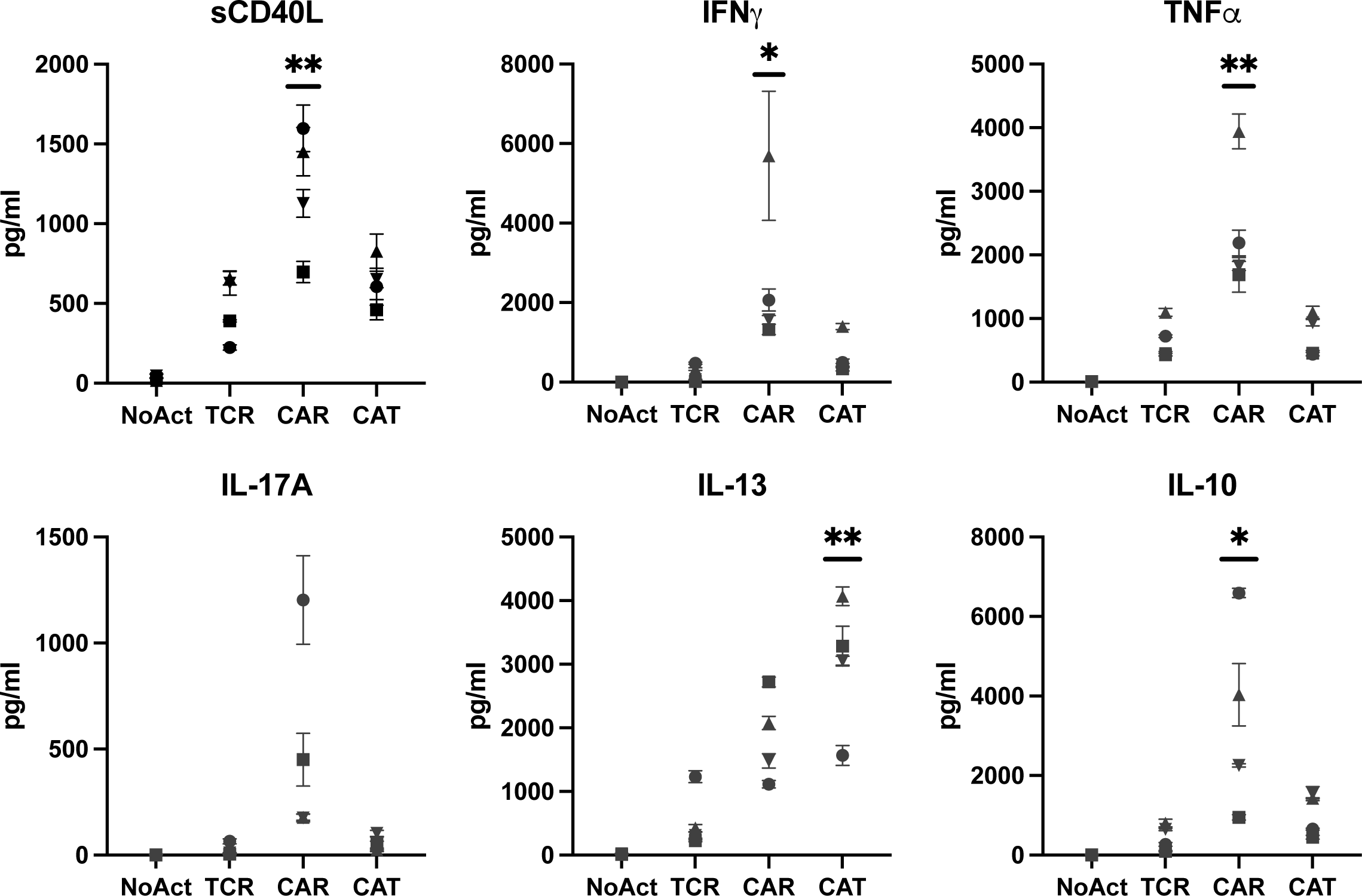
Low affinity CAR Tregs have dampened inflammatory cytokine secretion. Levels of cytokines secreted into the supernatant by No Act Tregs, TCR Tregs, CAR Tregs, and CAT Tregs 48h post-activation. Values represent mean ± SD of technical triplicates per blood donor. One-way ANOVA test with Tukey’s multiple comparison correction. **, p < 0.01; *, p < 0.05; ns, not significant.

Next, we sought to explore whether measuring the levels of the surface markers CD40L and TIGIT could help identify pro-inflammatory CAR Tregs and how these levels were affected by the affinity of the CAR. We activated TCR, CAR, and CAT Tregs with the respective irradiated K562 cell lines overnight and, following a 5-hour treatment with brefeldin A, we performed surface staining for CD40L and TIGIT, and then intracellular staining for IFNγ. While CAR Tregs and CAT Tregs both had higher expression of CD40L than TCR/CD28-activated Tregs (**Figure 10A**), CAT Tregs had TIGIT levels almost as high as TCR/CD28-activated Tregs (**Figure 10B**). Co-expression analysis revealed that, while the majority of TCR Tregs and CAT Tregs were TIGIT^+^CD40L^low^ cells, CAR Tregs were mostly TIGIT negative, with 20% of the cells being TIGIT^−^CD40L^hi^ cells (**Figure 10C**). Across the 4 subpopulations of CD40L and TIGIT expression combinations, high expression of CD40L correlated with high IFNγ production, with 20% of CD40L^hi^ CAR Tregs producing IFNγ versus only 5% of CD40L^low^ CAR Tregs (**Figure 10D**). Hence, CD40L surface expression correlates with IFNγ production in Tregs. Still, IFNγ-producing TCR Tregs and CAT Tregs were significantly less abundant than IFNγ-producing CAR Tregs irrespective of CD40L expression (**Figure 10D**), indicating that there are additional differences between CD40L^hi^ high affinity CAR-activated Tregs and CD40L^hi^ TCR/CD28-activated or low affinity CAR-activated Tregs.

**Figure 10.**
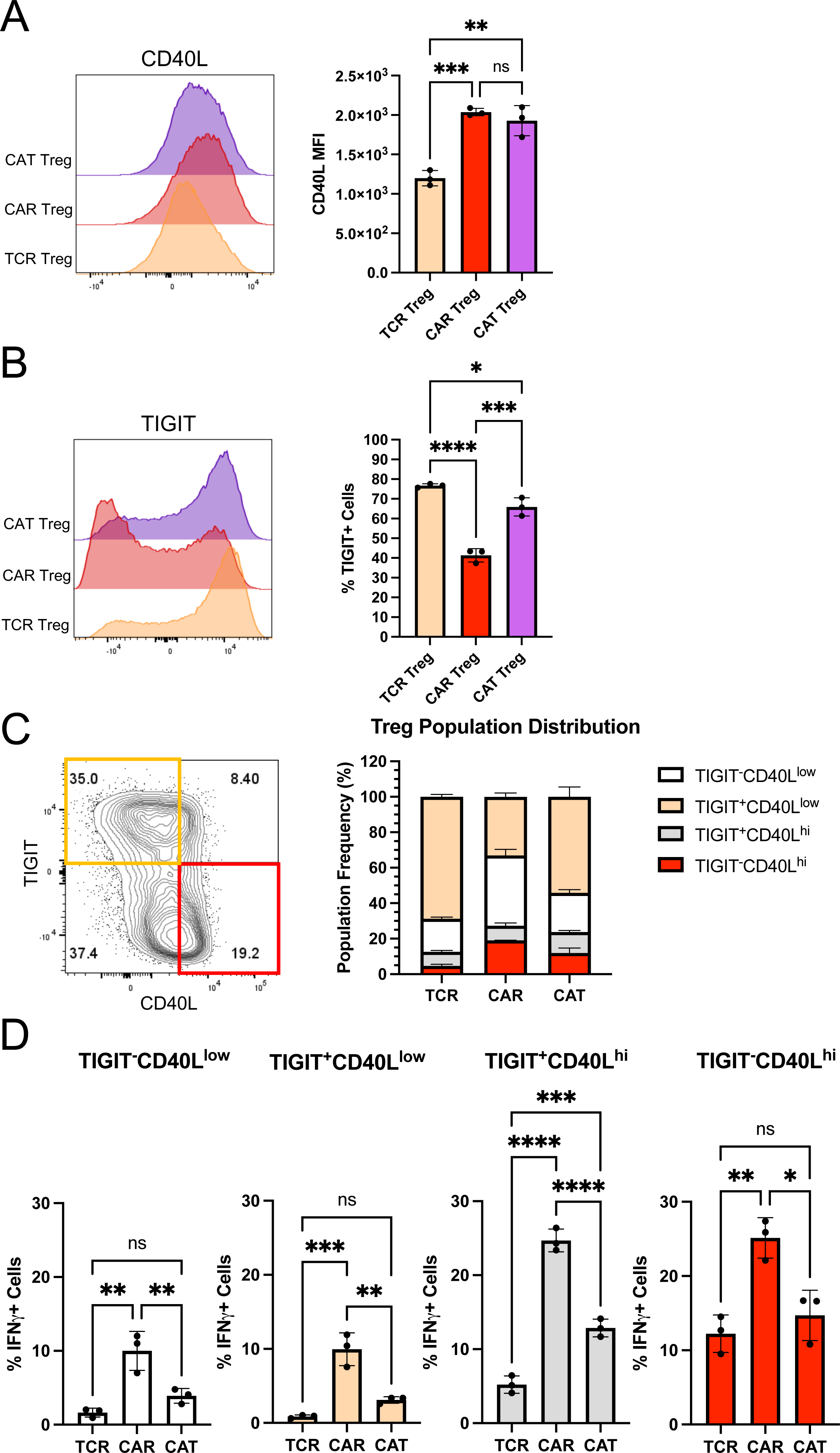
CD40L expression is associated with IFNγ production in CAR Tregs. (A) CD40L surface expression in TCR Tregs, CAR Tregs, and CAT Tregs 18h post-activation. Representative histograms on the left and summary data on the right. (B) TIGIT surface expression in TCR Tregs, CAR Tregs, and CAT Tregs 18h post-activation. Representative histograms on the left and summary data on the right. (C) Relative frequency of TIGIT^−^CD40L^low^, TIGIT^+^CD40L^low^, TIGIT^−^CD40L^hi^, and TIGIT^+^CD4L^hi^ cells among TCR Tregs, CAR Tregs, and CAT Tregs 18h post-activation. (D) Frequency of IFNG producing cells among TIGIT^−^CD40L^low^, TIGIT^+^CD40L^low^, TIGIT^−^CD40L^hi^, and TIGIT^+^CD4L^hi^ subpopulations for TCR Tregs, CAR Tregs, and CAT Tregs 18h post-activation. For Figures 10A, B, and D, values represent technical replicates of representative experiments. Bars represent mean ± SD. One-way ANOVA test with Tukey’s multiple comparison correction. ****, p < 0.0001; ***, p < 0.001; **, p < 0.01; *, p < 0.05; ns, not significant.

## DISCUSSION

The application of CAR technology to Tregs to induce or re-establish immune tolerance has been met with cautious optimism. While CAR engineered Tregs have shown promising results *in vitro* and in murine disease models of GvHD and skin graft rejection ^16–19^, their suboptimal efficacy in preclinical models of vascularized organ transplantation and autoimmune disease ^20,23,24^, settings where antigen-specific TCR Tregs have demonstrated efficacy ^26,75^, exposes the current limitations of CAR Treg-based strategies. This disparity underscores the need for a more complete understanding of how CAR Tregs function at a molecular level compared to their naturally activated (TCR/CD28) counterparts.

Unlike previous studies that relied on antibody- or antigen-coated beads for TCR activation ^76,77^, our study employed cellular targets for both CAR and TCR/CD28 activation with the goal of better mimicking physiological TCR and CAR synapses and their downstream signaling ^78^. In addition, we utilized a well-established CAR with a CD28-CD3zeta signaling domain with the goal of comparing CD28 and TCR/CD3 signaling delivered via a CAR and via the endogenous TCR and CD28 receptor. Our rationale for this comparative investigation is rooted in the fact that CAR constructs were originally designed and optimized for proinflammatory cytotoxic T cells. Consequently, we hypothesized that applying this same CAR architecture to immunosuppressive Tregs does not fully elicit or even disrupts Treg function, potentially jeopardizing their safe and effective clinical application.

On a first look, CAR and TCR/CD28-activated Tregs were similar in terms of activation marker upregulation, expansion, and stability (**Figure 2**). CAR Tregs, however, had lower CD25 levels across all donors (**Figure 2C**). This observation foreshadowed our findings that CAR Tregs were inferior at suppressing the proliferation of CD4^+^ T cells and CD8^+^ T cells (**Figures 3A and 3B**), an activity known to be dependent on IL-2 deprivation ^79^. CAR Tregs were also inferior at downregulating CD80 expression on target cells (**Figure 3C**), another important Treg suppression mechanism. Of note, CTLA4 was not differentially expressed between CAR Tregs and TCR Tregs, as determined by RNA-seq (**Table S5**). Interestingly, CAR Tregs were more cytotoxic towards target NALM6 cells (**Figure 3D**), a CD19-expressing B-cell leukemia, than TCR/CD28-activated Tregs. This could be due to the dramatic difference in affinity between a CAR scFv and a TCR. More specifically, the difference in cytotoxicity could be due to the CAR being slower at dissociating from its antigen than a TCR. The dissociation constant KD, which is inversely proportional to the binding affinity, of a TCR is normally in the range of 10^−4^ to 10^−7^ M ^41,72^. In contrast, the FMC63 CD19 CAR has a KD of 3.3 x 10^−10^ M and the CAT-13.1E10 CD19 CAR a KD of 1.4 x 10^−8^ M ^40^. The KD for a receptor is the ratio between how fast the receptor dissociates from its antigen, koff, and how fast the receptor binds to its antigen, kon. The koff for the FMC63 CD19 CAR is 6.8 x 10^−5^ s^−1^, whereas the koff for a TCR can vary from as fast as 10^−1^ s^−1^ to as slow as 10^−3^ s^−1^, which is still over 100 times faster than that of the FMC63 CAR ^40,72^. Seminal work showed that the longer a Treg is bound to a target dendritic cell, the more likely the Treg is to kill that cell ^52^. Strikingly, even if a TCR sequence is artificially mutated to generate a receptor with an affinity (KD) of 1.5 x 10^−8^ M, so a very similar KD to the low affinity CAR we tested in our work ^40^, the speed at which the CAR dissociates from its antigen is still lower than that of the TCR, with the koff for the CAT-131E10 CAR being 3.1 x 10^−3^ s^−1^ vs. the koff for the mutant TCR of 1.3 x 10^−3^ s^−1^ ^72^. Hence, the increased toxicity of CAR Tregs compared to TCR Tregs could be due to increased time bound to the target cells.

Of note, neither CAR Tregs nor TCR Tregs were cytotoxic towards CD19-expressing A549 cells (**Figure S1D**), engineered lung epithelial cancer cells, lending hope that CAR Tregs might not be directly cytotoxic towards non-immune tissues and organs. This possibility deserves special consideration, as CAR Tregs being currently tested in clinical trials (NCT05234190) target HLA-A2 expressed specifically in the transplanted organ to be protected from immune rejection ^29^.

Our functional assays suggested that CAR activation causes a shift from suppression to cytotoxicity (**Figure 3**). In line with this notion, CAR Tregs preferentially upregulated Teff cell inflammatory gene pathways (**Figure 4**, **Figure S2**) and uniquely produced inflammatory cytokines, notably IFNγ (**Figure 5**). IFNγ is an unwanted cytokine in the context of CAR Treg-based therapy, as it can lead to innate immune cell activation and HLA upregulation ^80^, thus being counterproductive in autoimmunity and organ transplant rejection. CAR Tregs did not, however, produce IL-2 (**Figure 5C**), cementing the idea that CAR Tregs remain stable Tregs upon activation. Lack of IL-2 production is a hallmark of Treg identity, with FOXP3 directly inhibiting transcription of the IL-2 gene ^81^. Curiously, IFNγ producing FOXP3^+^ Tregs have been previously described in autoimmunity and in solid tumors ^45,82^, suggesting that high affinity CAR activation may be tapping into Treg plasticity to elicit inflammatory cytokine production. CAR Teff cells also produced more IFNγ than TCR Teff cells (**Figure 5B**), suggesting that some aspect of high affinity CAR activation induces high IFNγ production across cell subsets. Previous reports have described the emergence of T helper-like Tregs that share transcription factor and chemokine gene expression patterns with T helper genes, e.g. Th1-like Tregs that express T-BET and CXCR3 ^83^. Yet, we did not find CAR activation to upregulate expression of TBX21, the gene coding for T-BET, in CAR Tregs at the bulk level, in spite of a 40-fold increase in IFNG expression (**Figure 6C**). Future profiling of gene expression at the single-cell level, as well as gene overexpression and deletion experiments, are poised to elucidate the gene circuitry conferring CAR Tregs partial Teff cell gene expression and exuberant cytokine and chemokine production.

Intriguingly, our study also identified heightened expression of CD40L in CAR Tregs (**Figure 6**), correlating with IFNγ expression (**Figure 10**). Activated CD4^+^ T helper cells express CD40L, which binds to CD40 on the surface of B cells; CD40L-CD40 signaling is required for high-titer high-affinity class-switched antibody production by B cells and for humoral memory formation ^64^. Tregs, in contrast, do not typically express CD40L, with CD40L negativity having been previously put forward as a strategy to isolate activated Tregs ^84,85^. While the implications of this *de novo* expression of CD40L in Tregs are not explored in the current study, they warrant further investigation, possibly including unwanted activation of CD40-expressing B cells and macrophages and concomitantly tissue damage ^86^. Of note, CD40L provides a potential surface marker to further purify and interrogate pro-inflammatory CAR Tregs in future studies.

Lowering CAR affinity by swapping the FMC63 scFv with the lower affinity CAT13.1E10 (CAT) scFv resulted in Tregs with a phenotype closer to that of TCR/CD28-activated Tregs, namely lower IFNγ production (**Figure 9**, **Figure 10**), higher TIGIT expression (**Figure 10**), and a lower frequency of CD40L-expressing cells (**Figure 10**). CAT Tregs also displayed higher suppression of CD4^+^ T cell proliferation, a greater downregulation of CD80 expression on target cells, and lower cytotoxicity towards NALM6 than CAR Tregs (**Figure 8**), establishing scFv affinity as a key parameter in CAR design for Tregs. Nevertheless, some differences between CAT Tregs and TCR Tregs subsisted, namely low expression of some chemokine receptor genes and higher secretion of some cytokines (**Figures 9, S5, and S6**).

The speed of translating Tregs to the clinic has been vertiginous, with only 10 years elapsing from their identification in humans in 2001 to their testing in graft-vs-host disease patients in 2011 ^5^. Yet, CAR Tregs are in their infancy as a strategy for immune regulation. Our work indicates that CAR Tregs can have a dual nature – pro-inflammatory yet still retaining key immunosuppressive features – calling for a more nuanced understanding of their complex signaling and functional outcomes if CAR Tregs are to become a safe and efficacious therapeutic modality. It also emphasizes how important it will be to tailor CAR constructs to Treg biology. Our data suggest that one possible avenue to achieve this is to ensure that the CAR affinity is not too high, lest it bestow Tregs with undesired inflammatory properties.

### Limitations of the study

While our study finds a clear unique phenotype in high affinity CAR-activated Tregs in comparison with TCR/CD28-activated Tregs and low affinity CAR-activated Tregs, only three CAR constructs specific for one target were used in this study. Further investigations are needed with different CAR constructs to cover a wider range of affinities, as well as a diversity of targets, as target molecule density on target cells has also been shown to influence CAR T-cell function ^87^. Moreover, some parameters of CAR constructs, such as the hinge and transmembrane domains ^87,88^, as well as alternative signaling domains ^50,89^, were not explored in the current study and may yield further insight. Another limitation of this study resides in the fact that it does not fully unveil the molecular mediators responsible for the induction of a pro-inflammatory phenotype and gene signature in Tregs by high affinity CAR activation. Finally, this study does not dissect the consequences of the unique CAR Treg phenotype discovered here *in vivo*, such as the effect of CAR Treg-derived IFNγ on a local milieu or the impact of CD40L-CD40 signaling on CAR Tregs and surrounding immune cells. Experiments using human CAR Tregs in humanized mouse models and murine CAR Tregs in immunocompetent mouse models can shed light on this aspect.

## METHODS

### Molecular Biology

CD64-2A-CD80, CD19ECD-PDGFRTM, and CD19CAR-2A-GFP lentiviral plasmids were synthesized by VectorBuilder Inc. (Chicago, IL). All genes were driven by an EF1A promoter. The CD19 CAR genes contained a CD8a signal peptide, an N-terminal Myc-tag, a single chain variable fragment (scFv) sequence recognizing human CD19, a CD8 hinge domain, a CD28 transmembrane domain, and a CD28-CD3zeta signaling domain. The high affinity anti-CD19 scFv sequence (FMC63) in the “CAR” CD19CAR construct was obtained from ^36^, the mutated CD28 signaling domain in the “PY3” CD19CAR construct was obtained from ^74^, and the low affinity anti-CD19 scFv sequence (CAT-13.1E10) in the “CAT” CD19CAR construct was obtained from ^40^. Lentivirus particles were produced by VectorBuilder Inc. and shipped to the laboratory, where they were stored in aliquots at −80°C until use. Construct sequences are available upon request.

### Regulatory T Cell Isolation

Human peripheral blood leukopaks from de-identified healthy donors were purchased from STEMCELL Technologies (Vancouver, Canada). CD4^+^ T cells and CD8^+^ T cells were enriched using the EasySep Human CD4^+^ T Cell Isolation Kit and EasySep Human CD8^+^ T Cell Isolation Kit (STEMCELL Technologies), respectively, as per manufacturer’s instructions. Enriched CD4^+^ T cells were then stained for CD4, CD25, and CD127, and CD4^+^CD25^hi^CD127^low^ regulatory T cells (Tregs), previously shown to be *bona fide* Tregs ^37,38^, and CD4^+^CD25^low^CD127^hi^ effector T (Teff) cells were purified by fluorescence-assisted cell sorting (FACS) using a BD FACS Aria II Cell Sorter (Beckton Dickinson, Franklin Lakes, NJ). Post-sort analyses confirmed greater than 99% purity. T cells were activated with anti-CD3/CD28 beads (Gibco, ThermoFisher Scientific) at a 1:1 ratio and recombinant human IL-2 (Peprotech, ThermoFisher Scientific), and expanded in RPMI 1640 medium supplemented with 10% fetal bovine serum (FBS), glutamax, penicillin-streptomycin, HEPES, non-essential amino acids (NEAA), and sodium pyruvate (all from Gibco, ThermoFisher Scientific). Tregs were cultured with 1,000 IU/ml IL-2, CD4^+^ Teff cells with 100 IU/ml IL-2, and CD8^+^ T cells with 300 IU/ml IL-2 ^48^. Antibodies used for FACS and flow cytometry can be found in **Table S9**.

### T Cell Transduction and Expansion

Two days after activation, T cells were transduced with CAR lentivirus at a multiplicity of infection (MOI) of 1 (1 particle per cell) in the presence of IL-2. After adding the lentivirus, T cells were centrifuged at 1,000 g at 32°C for 1 hour. Following transduction, T cells were maintained and expanded in RPMI10 medium with fresh medium and IL-2 being given every two days. CAR-expressing T cells were FACS sorted based on reporter GFP expression.

### CAR Treg Activation, Stability, and Expansion

CAR Tregs were co-cultured with irradiated K562 (No Activation), CD19-expressing K562 (CAR Activation) or CD64- and CD80-expressing K562 previously loaded with anti-CD3 antibody (OKT3, Biolegend, San Diego, CA) at 1 μg/ml for 1 hour ^39^ (TCR/CD28 Activation) at a 1:1 ratio of CAR Tregs to K562 cells in RPMI10 medium supplemented with 1,000 IU/ml IL-2. Surface expression of CD71 and CD25 (Activation) was assessed at 48 hours by flow cytometry. Parallel co-cultures were kept for one week to assess expression of FOXP3 and HELIOS (Stability) by intracellular staining using the FOXP3/Transcription Factor Staining Buffer Set (eBioscience, ThermoFisher Scientific), according to manufacturer’s instructions. Cell numbers were also assessed at this time (Expansion). Flow cytometry data was acquired in a 5-laser Beckman Coulter CytoFLEX flow cytometer or a 3-laser Cytek Northern Lights spectral flow cytometer. FlowJo v10.9 software (BD Life Sciences, Franklin Lakes, NJ) was used for flow cytometry data analysis.

### T Cell Suppression Assay

CAR Tregs were activated via CAR (with irradiated CD19-K562 cells), via TCR/CD28 (with irradiated CD64-CD80-K562 cells loaded with anti-CD3 OKT3 antibody) or left resting (with irradiated K562 cells) at a 1:1 Treg to target cell ratio in round bottom 96-well plates. In parallel, CD4^+^ and CD8^+^ T responder (Tresp) cells were mixed at a 1:1 ratio, labeled with CellTrace Violet (CTV) or CellTrace Far Red (CTFR) according to the manufacturer’s instructions (Invitrogen, ThermoFisher Scientific), and activated with anti-CD3/CD28 beads at a 1:2 bead to cell ratio overnight ^48,90^. The following day, Tresp cells were debeaded and co-incubated with activated Tregs in round bottom 96-well plates at different Treg:Tresp ratios for three days in the absence of exogenous IL-2 ^48,90^. Co-cultures were then harvested, stained for CD4 and CD8, and CTV or CTFR dye dilution measured via flow cytometry.

### Artificial Antigen Presenting Cell Suppression Assay

CAR^+^ Tregs were incubated with NALM6 cells and CAR^−^ Tregs were incubated with CD64-CD80-NALM6 loaded with anti-CD3 for 4 days. Co-cultures were then harvested and CD80 surface expression assessed using flow cytometry.

### Monocyte Isolation and Dendritic Cell Differentiation

Human CD14^+^ monocytes were isolated from leukopaks using the EasySep Human CD14^+^ Positive Selection Kit (STEMCELL Technologies) and differentiated into monocyte-derived dendritic cells (moDCs) using the ImmunoCult Dendritic Cell Culture Kit (STEMCELL Technologies), as per manufacturer’s instructions. Complete moDC maturation was assessed by surface expression of CD11c, CD80, CD83, and CD86 using flow cytometry. Cells were frozen in freezing medium (90% FBS, 10% DMSO) and stored in liquid nitrogen until being thawed for assays.

### Dendritic Cell Suppression Assay

Monocyte-derived dendritic cells (moDCs) were thawed on the day of the experiment and plated in each well supplemented with 50 ng/mL IFNγ (STEMCELL Technologies) for overnight activation. In parallel, Tregs were activated via CAR (with irradiated CD19-K562 cells), via TCR/CD28 (with irradiated CD64-CD80-K562 cells loaded with anti-CD3 OKT3 antibody) or left resting (with irradiated K562 cells) at a 1:1 Treg to target cell ratio. The next day, IFNγ was washed off from moDCs, then Tregs were co-cultured with moDCs for 3 days. Co-cultures were then harvested and stained with CD4, CD11c, CD80, CD83, and CD86. Suppression of moDC was gauged based on the surface expression level of CD80 and CD86, as assessed by flow cytometry ^50^.

### Cytotoxicity assay

CAR^+^ Tregs were incubated with NALM6 cells and CAR^−^ Tregs were incubated with CD64-CD80-NALM6 loaded with anti-CD3 (OKT3 antibody) for 24h. Target cell killing was then assessed using the CyQUANT Cytotoxicity Lactate Dehydrogenase (LDH) Release (a measure of cell death) Assay kit (Thermofisher Scientific) as per manufacturer’s instructions.

### CRISPR/Cas9 gene editing

Two days after activation with anti-CD3/28 beads and 1,000 IU/ml IL-2, Tregs were debeaded and electroporated with Cas9 (TrueCut v2, ThermoFisher Scientific) and guide RNA (Synthego, Redwood City, CA) ribonucleoprotein complexes (RNP) using a Neon system (ThermoFisher Scientific) with settings 2200 V, 20 ms, 1 pulse. Electroporated cells were recovered in antibiotic-free RPMI10 with IL-2 and expanded until analysis. The guide RNA sequence used to target the PRF1 gene (encoding the perforin protein) was 5’-CCTTCCCAGTGGACACACAA-3’. Control wild-type (WT) cells were electroporated with Cas9 alone. CRISPR/Cas9 genome editing efficiency was assessed by PCR amplification of a 500 bp region of the genomic DNA containing the PRF1 gRNA cutting site, using the forward primer 5’-AAGGGAGCAGTCATCCTCCA-3’ and the reverse primer 5’-CATTGCTGGTGGGCTTAGGA-3’, followed by Sanger sequencing (Eurofins Genomics, Louisville, KY) and sequence analysis using Tracking of Indels by Decomposition (TIDE, https://tide.nki.nl/) to obtain indel frequency ^56^.

### Whole Transcriptome RNA-seq Analysis

CAR Tregs and CAR Teff cells were co-cultured with irradiated K562 (No Activation), CD19-K562 (CAR Activation) or CD64-CD80-K562 loaded with anti-CD3 antibody (TCR/CD28 Activation) at a 1:1 ratio in RPMI10 medium. CAR Treg co-cultures were supplemented with 1,000 IU/ml IL-2. After 24h, CD4^+^ cells were isolated using the EasySep Human CD4 Positive Selection Kit (STEMCELL Technologies), following the manufacturer’s instructions. RNA-seq libraries were built using poly-A selection and paired-end sequencing was performed with the Illumina NovaSeq 6000 platform. For data analysis, FastQC was first applied to assess the quality of raw sequencing reads. Alignment was then performed with STAR (Spliced Transcripts Alignment to a Reference) alignment software ^91^ using the most recent build of the human GENCODE reference genome (Release 44, GRCh38.p14). Next, Samtools were employed for filtering and sorting uniquely aligned reads and FeatureCounts for annotating and quantifying raw gene counts ^92^. Gene transfer format files for gene annotation from GENCODE (hg38/GRCh38) were then obtained. DESeq2 ^93^ was used for normalization and downstream differential gene expression analysis. Genes showing a false discovery rate (FDR) < 0.05 and absolute log2 fold change > 1 in magnitude were considered differentially expressed in pair-wise comparisons. The topmost significantly differentially upregulated genes were used for gene set enrichment analysis (GSEA) ^62^. Some RNA-seq data inspection and visualization was performed with the help of Venny 2.0 (https://bioinfogp.cnb.csic.es/tools/venny/index2.0.2.html) and iDEP 2.0 ^94^ (http://bioinformatics.sdstate.edu/idep/). Raw and processed data to support the findings of this study have been deposited in GEO under accession number: xxx. Code used to analyze the RNA-seq data in this paper can be found at xxx.

### Cytokine Secretion

Supernatants from Treg and Teff cell co-cultures with K562 target cell lines were collected, stored at −0°C and shipped to EveTech Inc. (Calgary, Canada) for cytokine quantitation using multiplex ELISA.

### Intracellular Cytokine Production

CAR Tregs and CAR Teff cells were activated overnight via CAR (with irradiated CD19-K562 cells), via TCR/CD28 (with irradiated CD64-CD80-K562 cells loaded with anti-CD3 antibody) or left resting (with irradiated K562 cells) at a 1:1 Treg to target cell ratio in round bottom 96-well plates. The following day, co-cultures were treated with Brefeldin A (Biolegend) for 5h and harvested for intracellular cytokine staining with the FOXP3/Transcription Factor Staining Buffer Set (eBioscience, ThermoFisher Scientific), according to manufacturer’s instructions.

### Quantitative polymerase chain reaction (qPCR)

Total RNA from CAR and TCR/CD28 activated Tregs 24h post-activation was isolated using Trizol (ThermoFisher Scientific), according to manufacturer’s instructions. A total of 1000 ng of RNA was used for cDNA synthesis with the High-Capacity cDNA Reverse Transcription Kit (Bio-Rad, Hercules, CA). Real-time PCR was performed with iTaq Universal SYBR Green Supermix (Bio-Rad) on a Bio-Rad Real-time System C1000 Thermal Cycler. Target gene Ct values were normalized to RPL13A Ct value. Sequences of the primers used for qPCR can be found in **Table S10**.

### Statistics

Statistical analyses were performed using GraphPad Prism v10.0.0 (GraphPad Software, La Jolla, CA).

## SUPPLEMENTAL INFORMATION

This manuscript contains 6 supplemental figures and 10 supplemental tables.

## ACKNOWLEDGMENTS

This work was supported by Medical University of South Carolina and Hollings Cancer Center startup funds, Human Islet Research Network (HIRN) Emerging Leader in Type 1 Diabetes grant U24DK104162-07, American Cancer Society (ACS) Institutional Research Grant IRG-19-137-20, Diabetes Research Connection (DRC) grant IPF 22-1224, and Swim Across America (SAA) grant 23-1579 to L.M.R.F., and Cellular, Biochemical and Molecular Sciences training grant 5T32GM132055 to R.W.C. Supported in part by the Flow Cytometry and Cell Sorting Shared Resource, Hollings Cancer Center, Medical University of South Carolina (P30 CA138313). We thank past and present members of the Ferreira Lab for helpful discussions.

## AUTHOR CONTRIBUTIONS

L.M.R.F. and R.W.C. conceived the study. L.M.R.F. supervised the study. L.M.R.F., R.W.C, R.A.R., E.A., S.V., and M.J.R. performed experiments. L.M.R.F., R.W.C., R.A.R., B.G., E.A., A.A.C., and S.B. performed data analysis. L.M.R.F. and R.W.C. wrote the manuscript. All authors reviewed and approved the manuscript.

## DECLARATION OF INTERESTS

L.M.R.F. is the inventor on a provisional patent based on results from this work, an inventor on provisional and licensed patents on engineered immune cells, and a consultant with Guidepoint Global. The other authors declare no conflicts of interest.

## SUPPLEMENTAL FIGURE LEGENDS

**Figure S1.**
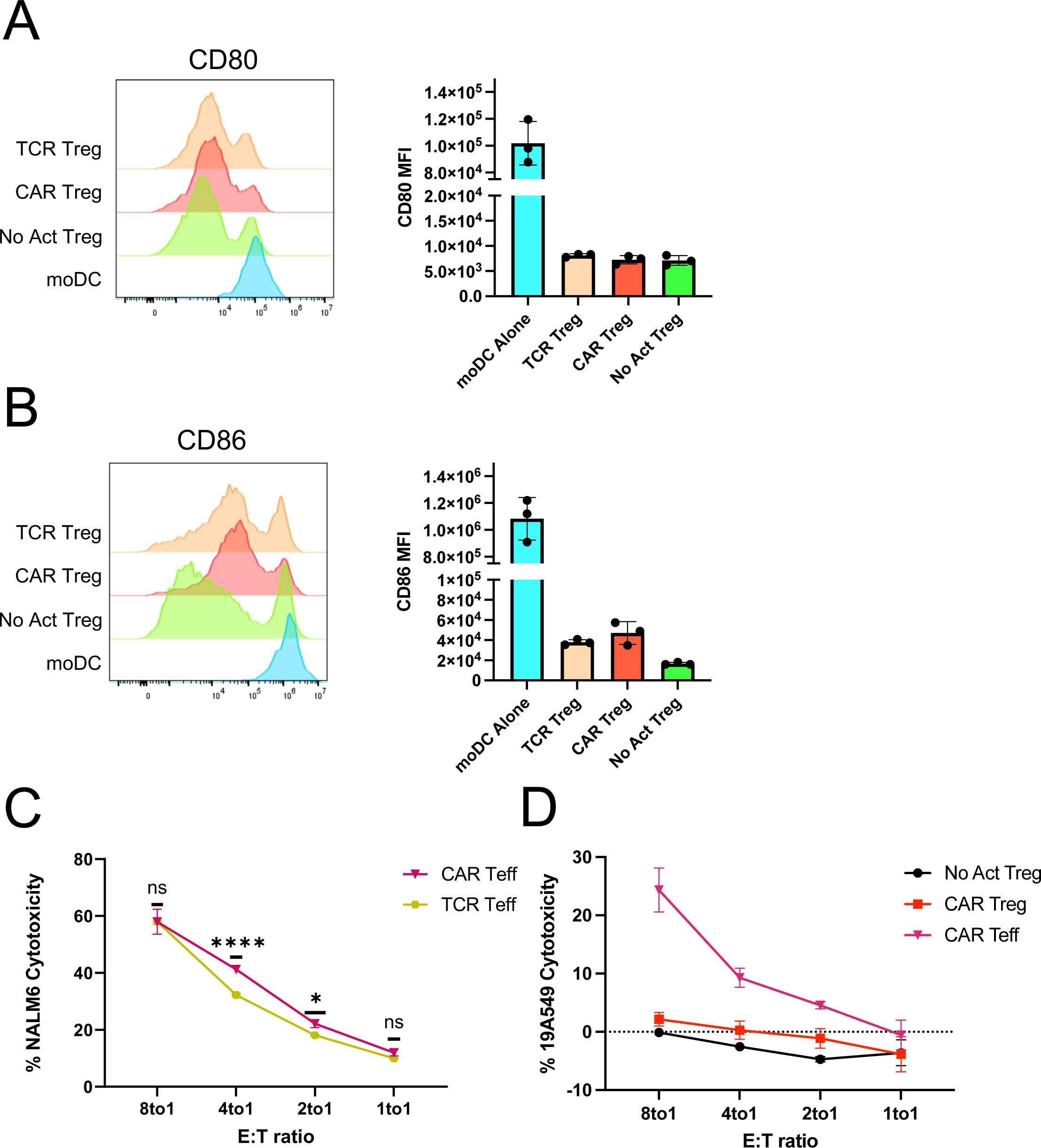
CAR Tregs are not cytotoxic towards epithelial cells. (A) CD80 surface expression on monocyte-derived dendritic cells (moDCs) 4 days after co-incubation with No Act Tregs, TCR Tregs or CAR Tregs. Representative histogram on the left and summary data on the right. (B) CD86 surface expression on moDCs 4 days after co-incubation with No Act Tregs, TCR Tregs or CAR Tregs. Representative histogram on the left and summary data on the right. (C) Teff cytotoxicity towards target NALM6 cells at different effector to target (E:T) ratios. (D) Treg and Teff cytotoxicity towards target CD19-A549 cells at different E:T ratios.

**Figure S2.**
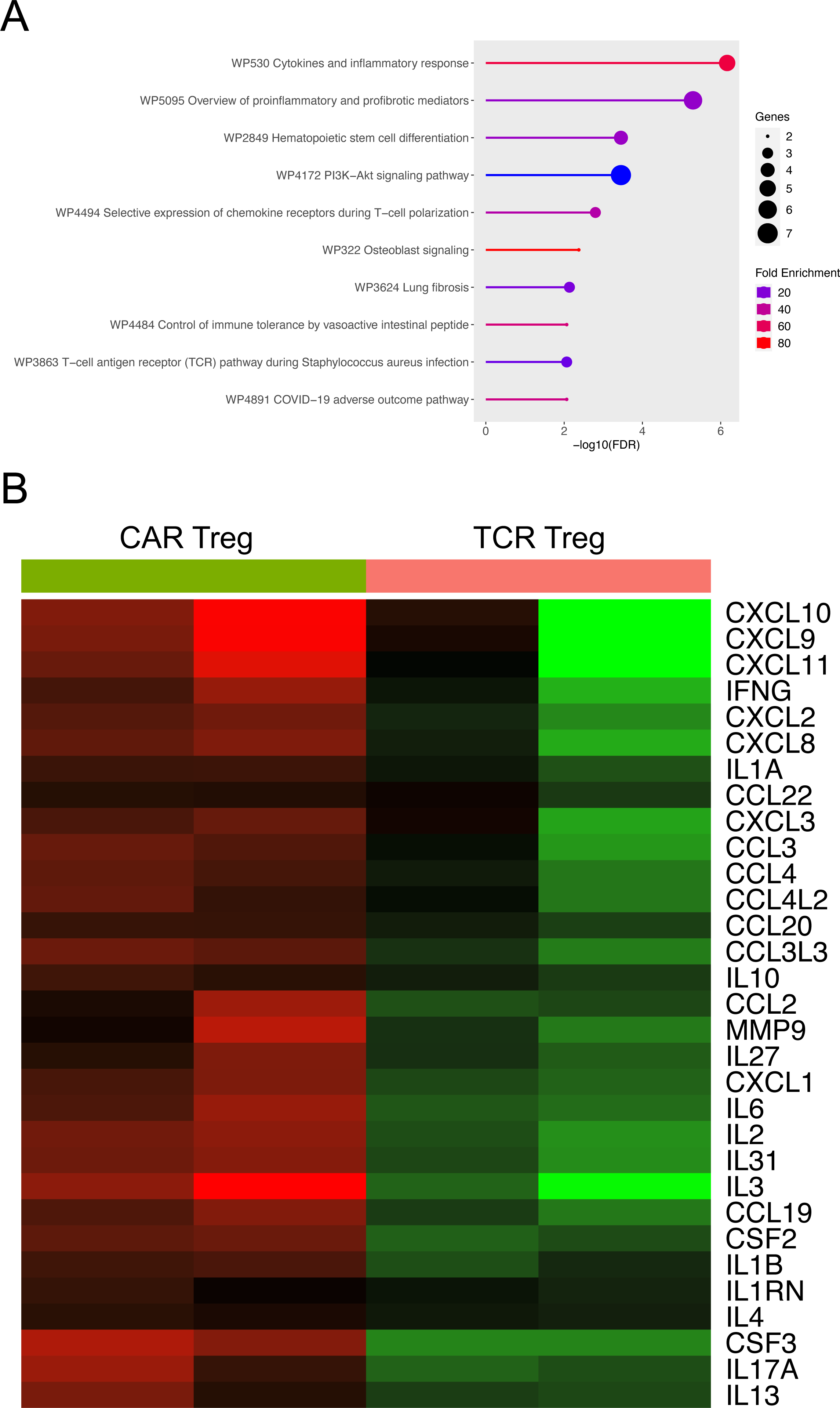
Inflammatory gene and gene pathways upregulated by CAR activation in Tregs. (A) WikiPathways gene set enrichment analysis (GSEA) of CAR Tregs vs. TCR Tregs. FDR, false discovery rate. (B) Heatmap of CAR Treg and TCR Treg proinflammatory and profibrotic mediator gene expression, as determined by RNA-seq.

**Figure S3.**
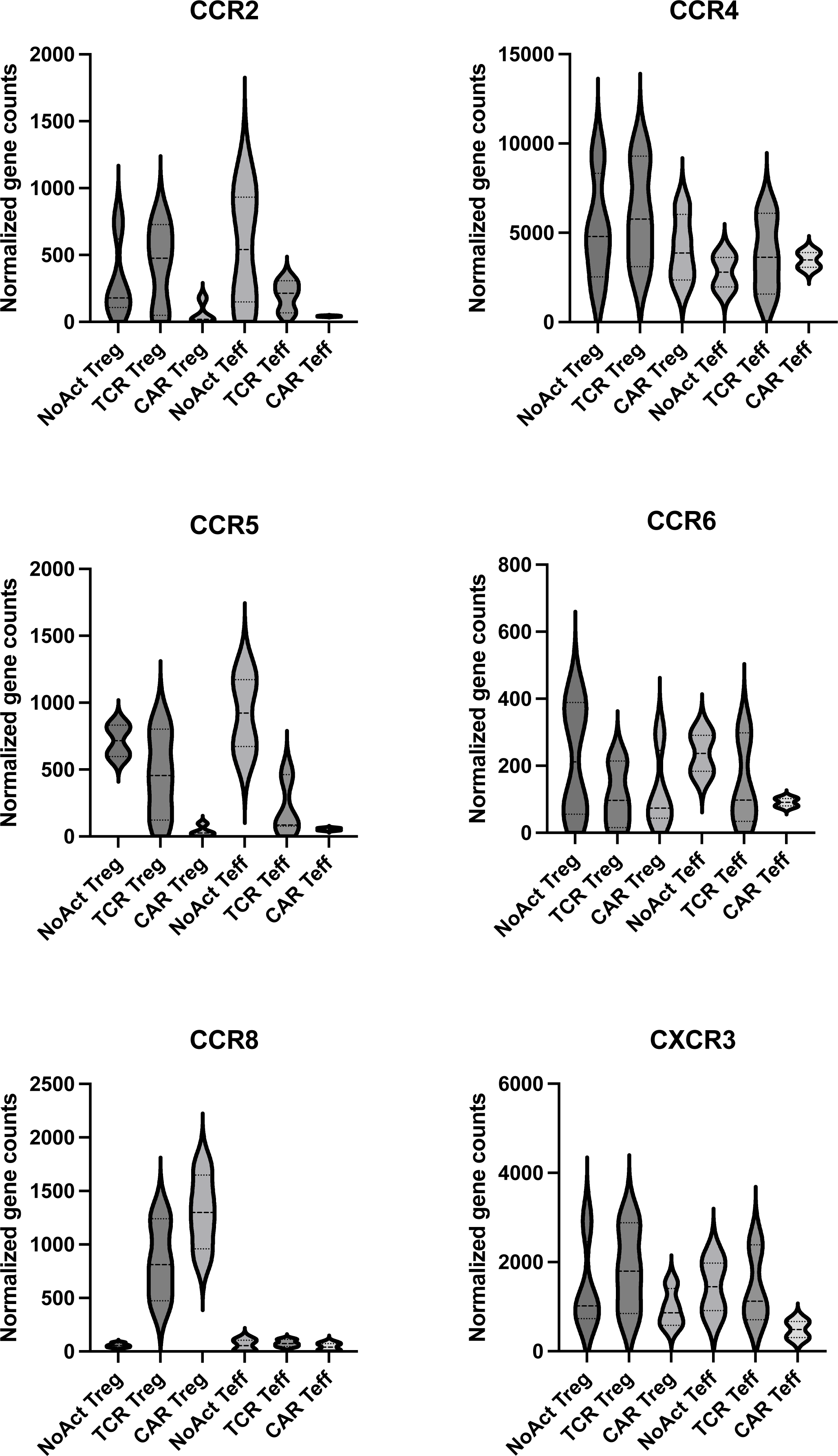
Chemokine receptor gene expression levels in No Act Tregs, TCR Tregs, CAR Tregs, No Act Teff, TCR Teff, and CAR Teff. Violins represent mean ± SD of RNA-seq values from different blood donors.

**Figure S4.**
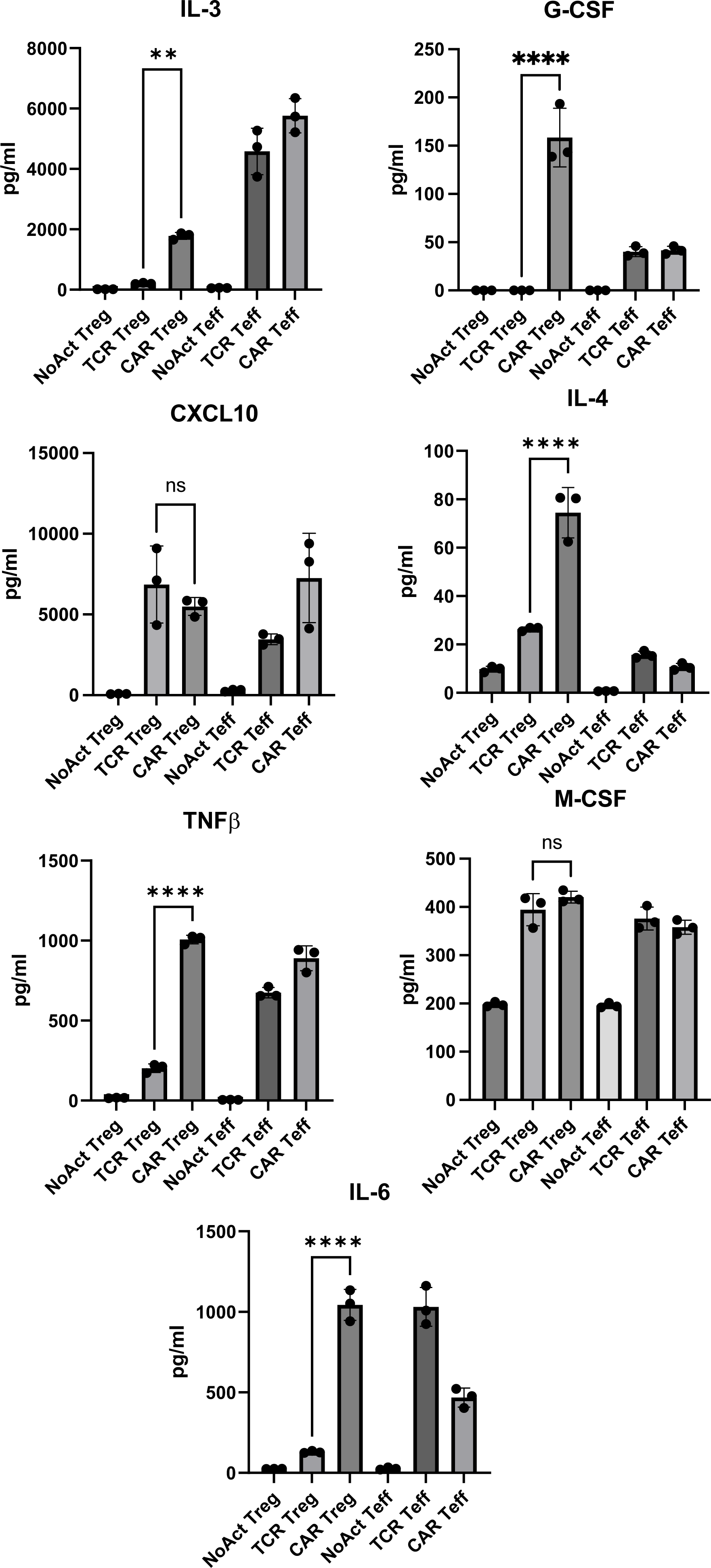
Cytokine secretion levels by No Act Tregs, TCR Tregs, CAR Tregs, No Act Teff, TCR Teff, and CAR Teff. Values represent technical replicates of representative experiments. Bars represent mean ± SD. One-way ANOVA test with Tukey’s multiple comparison correction. ****, p < 0.0001; ***, p < 0.001; **, p < 0.01; *, p < 0.05; ns, not significant.

**Figure S5.**
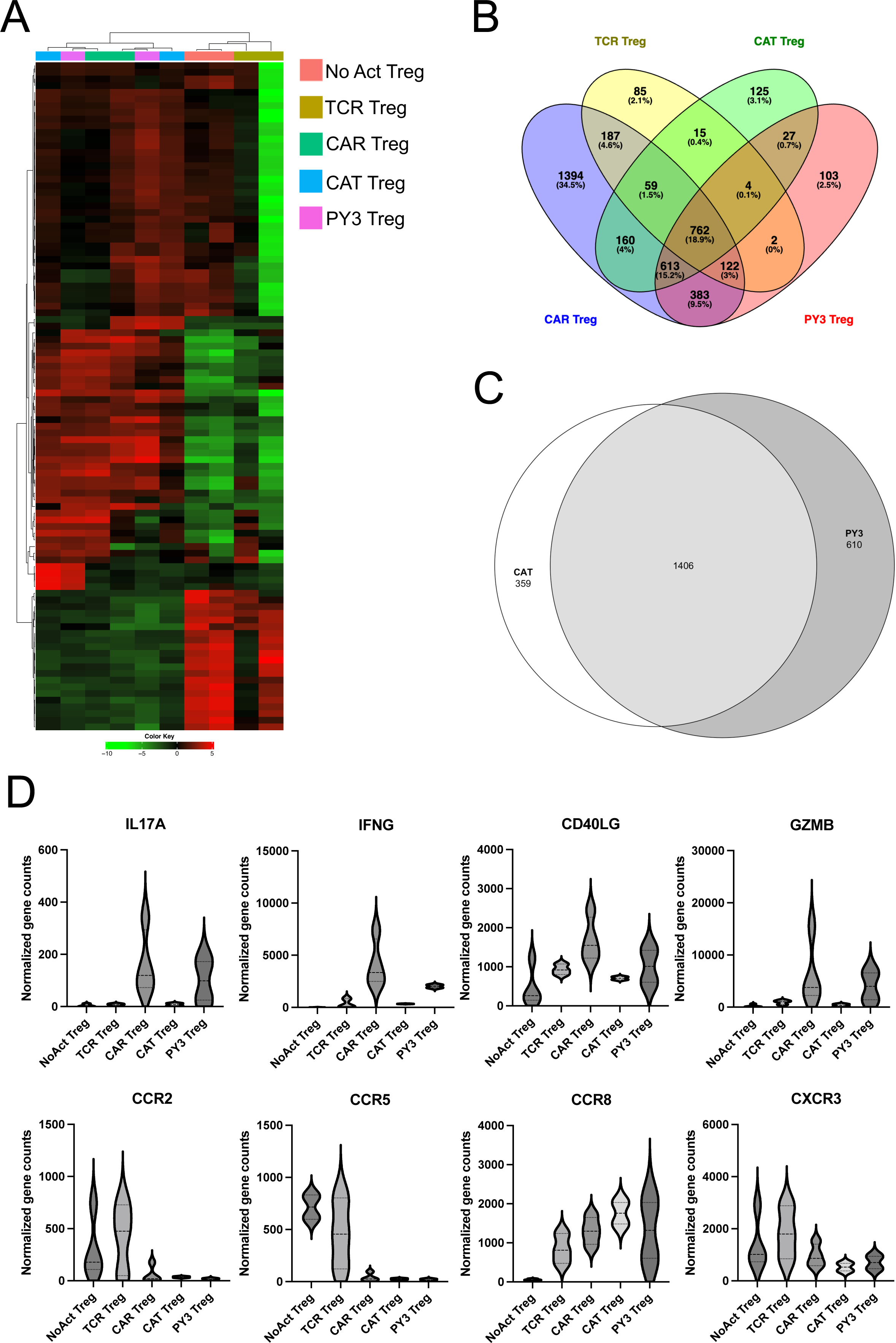
CAT Tregs have lower inflammatory gene expression levels than CAR Tregs. (A) Heatmap clustered by column (sample) and by row (gene) with top 100 most differentially expressed genes between No Act Tregs, TCR Tregs, CAR Tregs, CAT Tregs, and PY3 Tregs. (B) Venn diagram with genes upregulated in TCR Tregs, CAR Tregs, CAT Tregs, and PY3 Tregs in relation to their respective No Act cell types. Number of genes and respective percentage of the total number of genes are indicated in each intersection. (C) Venn diagram with genes upregulated in CAT Tregs and in PY3 Tregs. (D) Inflammatory, cytotoxic, and chemokine receptor gene expression levels in No Act Tregs, TCR Tregs, CAR Tregs, CAT Tregs, and PY3 Tregs. Violins represent mean ± SD of of RNA-seq values from different blood donors.

**Figure S6.**
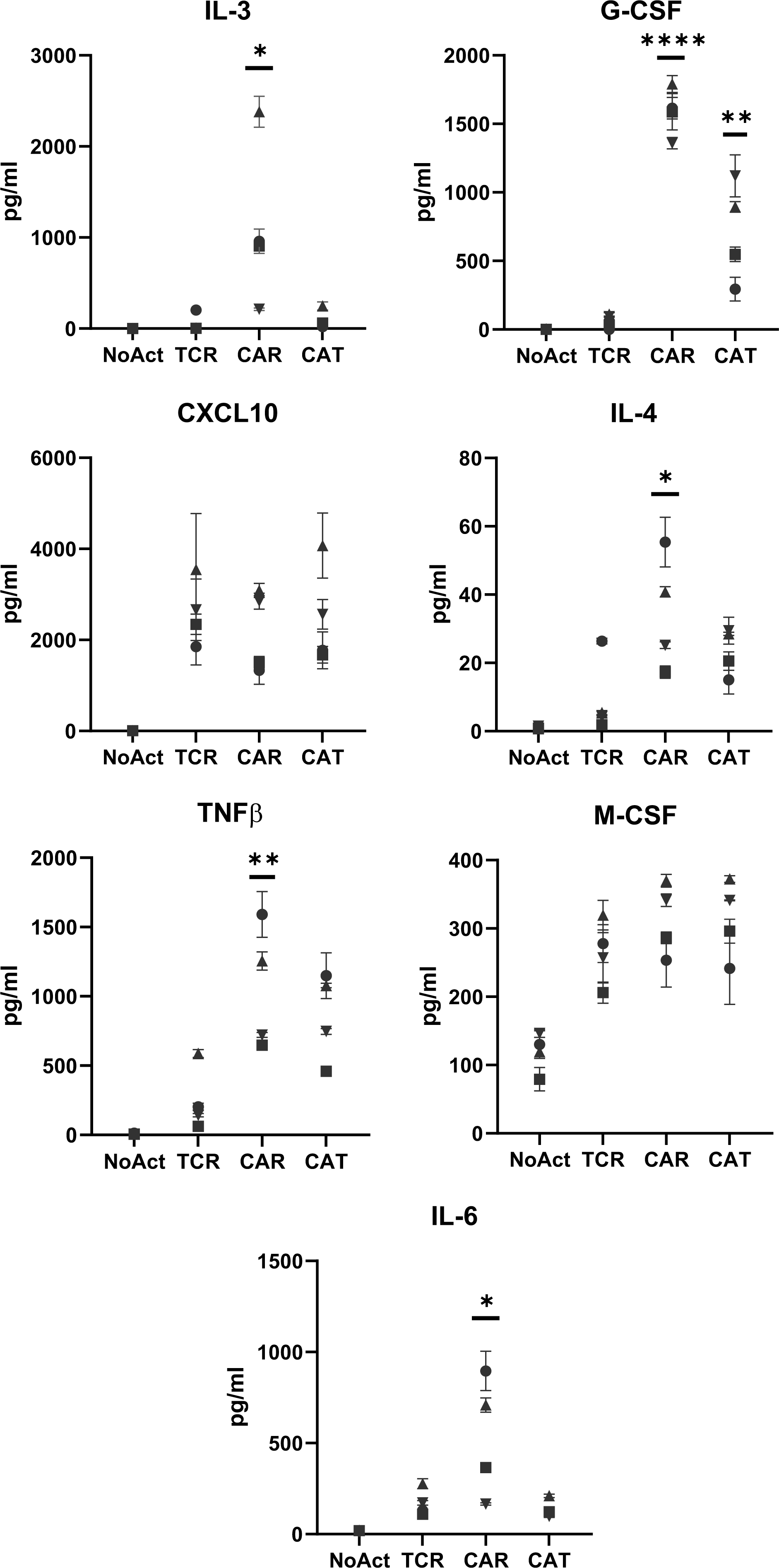
Cytokine secretion levels by No Act Tregs, TCR Tregs, CAR Tregs, and CAT Tregs. Values are the mean ± SD of technical triplicates per blood donor. One-way ANOVA test with Tukey’s multiple comparison correction. ****, p < 0.0001; ***, p < 0.001; **, p < 0.01; *, p < 0.05; ns, not significant.

## SUPPLEMENTAL TABLES

**Table S1. Differentially expressed genes in CAR Tregs compared with NoAct Tregs.**

**Table S2. Differentially expressed genes in TCR Tregs compared with NoAct Tregs.**

**Table S3. Differentially expressed genes in CAR Teff compared with NoAct Teff.**

**Table S4. Differentially expressed genes in TCR Teff compared with NoAct Teff.**

**Table S5. Differentially expressed genes in CAR Tregs compared with TCR Tregs.**

**Table S6. Genes upregulated in CAR Tregs, CAR Teff, and TCR Teff, but not in TCR Tregs.**

**Table S7. Genes upregulated only in TCR Tregs and not in CAR Tregs, CAR Teff or TCR Teff.**

**Table S8. Genes upregulated in PY3 Tregs and not in CAT Tregs.**

**Table S9. Flow cytometry antibodies and dyes used in this study.**

**Table S10. Primers used in this study.**

## REFERENCES

1. Ettenger, R., Chin, H., Kesler, K., Bridges, N., Grimm, P., Reed, E.F., Sarwal, M., Sibley, R., Tsai, E., Warshaw, B., and Kirk, A.D. (2017). Relationship Among Viremia/Viral Infection, Alloimmunity, and Nutritional Parameters in the First Year After Pediatric Kidney Transplantation. Am J Transplant 17, 1549–1562. 10.1111/ajt.14169.

2. Nelson, J., Alvey, N., Bowman, L., Schulte, J., Segovia, M.C., McDermott, J., Te, H.S., Kapila, N., Levine, D.J., Gottlieb, R.L., et al. (2022). Consensus recommendations for use of maintenance immunosuppression in solid organ transplantation: Endorsed by the American College of Clinical Pharmacy, American Society of Transplantation, and the International Society for Heart and Lung Transplantation. Pharmacotherapy 42, 599–633. 10.1002/phar.2716.

3. Shivaswamy, V., Boerner, B., and Larsen, J. (2016). Post-Transplant Diabetes Mellitus: Causes, Treatment, and Impact on Outcomes. Endocr Rev 37, 37–61. 10.1210/er.2015-1084.

4. Ghobadinezhad, F., Ebrahimi, N., Mozaffari, F., Moradi, N., Beiranvand, S., Pournazari, M., Rezaei-Tazangi, F., Khorram, R., Afshinpour, M., Robino, R.A., et al. (2022). The emerging role of regulatory cell-based therapy in autoimmune disease. Front Immunol 13, 1075813. 10.3389/fimmu.2022.1075813.

5. Ferreira, L.M.R., Muller, Y.D., Bluestone, J.A., and Tang, Q. (2019). Next-generation regulatory T cell therapy. Nat Rev Drug Discov 18, 749–769. 10.1038/s41573-019-0041-4.

6. Hori, S., Nomura, T., and Sakaguchi, S. (2003). Control of regulatory T cell development by the transcription factor Foxp3. Science 299, 1057–1061. 10.1126/science.1079490.

7. Fontenot, J.D., Gavin, M.A., and Rudensky, A.Y. (2003). Foxp3 programs the development and function of CD4+CD25+ regulatory T cells. Nat Immunol 4, 330–336. 10.1038/ni904.

8. Khattri, R., Cox, T., Yasayko, S.A., and Ramsdell, F. (2003). An essential role for Scurfin in CD4+CD25+ T regulatory cells. Nat Immunol 4, 337–342. 10.1038/ni909.

9. Tay, C., Tanaka, A., and Sakaguchi, S. (2023). Tumor-infiltrating regulatory T cells as targets of cancer immunotherapy. Cancer Cell 41, 450–465. 10.1016/j.ccell.2023.02.014.

10. Li, J., Tan, J., Martino, M.M., and Lui, K.O. (2018). Regulatory T-Cells: Potential Regulator of Tissue Repair and Regeneration. Front Immunol 9, 585. 10.3389/fimmu.2018.00585.

11. Bluestone, J.A., Buckner, J.H., Fitch, M., Gitelman, S.E., Gupta, S., Hellerstein, M.K., Herold, K.C., Lares, A., Lee, M.R., Li, K., et al. (2015). Type 1 diabetes immunotherapy using polyclonal regulatory T cells. Sci Transl Med 7, 315ra189. 10.1126/scitranslmed.aad4134.

12. Tang, Q., Leung, J., Peng, Y., Sanchez-Fueyo, A., Lozano, J.J., Lam, A., Lee, K., Greenland, J.R., Hellerstein, M., Fitch, M., et al. (2022). Selective decrease of donor-reactive T(regs) after liver transplantation limits T(reg) therapy for promoting allograft tolerance in humans. Sci Transl Med 14, eabo2628. 10.1126/scitranslmed.abo2628.

13. Sawitzki, B., Harden, P.N., Reinke, P., Moreau, A., Hutchinson, J.A., Game, D.S., Tang, Q., Guinan, E.C., Battaglia, M., Burlingham, W.J., et al. (2020). Regulatory cell therapy in kidney transplantation (The ONE Study): a harmonised design and analysis of seven non-randomised, single-arm, phase 1/2A trials. Lancet 395, 1627–1639. 10.1016/S0140-6736(20)30167-7.

14. June, C.H., and Sadelain, M. (2018). Chimeric Antigen Receptor Therapy. N Engl J Med 379, 64–73. 10.1056/NEJMra1706169.

15. Cappell, K.M., and Kochenderfer, J.N. (2023). Long-term outcomes following CAR T cell therapy: what we know so far. Nat Rev Clin Oncol 20, 359–371. 10.1038/s41571-023-00754-1.

16. Muller, Y.D., Ferreira, L.M.R., Ronin, E., Ho, P., Nguyen, V., Faleo, G., Zhou, Y., Lee, K., Leung, K.K., Skartsis, N., et al. (2021). Precision Engineering of an Anti-HLA-A2 Chimeric Antigen Receptor in Regulatory T Cells for Transplant Immune Tolerance. Front Immunol 12, 686439. 10.3389/fimmu.2021.686439.

17. MacDonald, K.G., Hoeppli, R.E., Huang, Q., Gillies, J., Luciani, D.S., Orban, P.C., Broady, R., and Levings, M.K. (2016). Alloantigen-specific regulatory T cells generated with a chimeric antigen receptor. J Clin Invest 126, 1413–1424. 10.1172/JCI82771.

18. Noyan, F., Zimmermann, K., Hardtke-Wolenski, M., Knoefel, A., Schulde, E., Geffers, R., Hust, M., Huehn, J., Galla, M., Morgan, M., et al. (2017). Prevention of Allograft Rejection by Use of Regulatory T Cells With an MHC-Specific Chimeric Antigen Receptor. Am J Transplant 17, 917–930. 10.1111/ajt.14175.

19. Boardman, D.A., Philippeos, C., Fruhwirth, G.O., Ibrahim, M.A., Hannen, R.F., Cooper, D., Marelli-Berg, F.M., Watt, F.M., Lechler, R.I., Maher, J., et al. (2017). Expression of a Chimeric Antigen Receptor Specific for Donor HLA Class I Enhances the Potency of Human Regulatory T Cells in Preventing Human Skin Transplant Rejection. Am J Transplant 17, 931–943. 10.1111/ajt.14185.

20. Wagner, J.C., Ronin, E., Ho, P., Peng, Y., and Tang, Q. (2022). Anti-HLA-A2-CAR Tregs prolong vascularized mouse heterotopic heart allograft survival. Am J Transplant. 10.1111/ajt.17063.

21. Ellis, G.I., Coker, K.E., Winn, D.W., Deng, M.Z., Shukla, D., Bhoj, V., Milone, M.C., Wang, W., Liu, C., Naji, A., et al. (2022). Trafficking and persistence of alloantigen-specific chimeric antigen receptor regulatory T cells in Cynomolgus macaque. Cell Rep Med 3, 100614. 10.1016/j.xcrm.2022.100614.

22. Tenspolde, M., Zimmermann, K., Weber, L.C., Hapke, M., Lieber, M., Dywicki, J., Frenzel, A., Hust, M., Galla, M., Buitrago-Molina, L.E., et al. (2019). Regulatory T cells engineered with a novel insulin-specific chimeric antigen receptor as a candidate immunotherapy for type 1 diabetes. J Autoimmun 103, 102289. 10.1016/j.jaut.2019.05.017.

23. Obarorakpor, N., Patel, D., Boyarov, R., Amarsaikhan, N., Cepeda, J.R., Eastes, D., Robertson, S., Johnson, T., Yang, K., Tang, Q., and Zhang, L. (2023). Regulatory T cells targeting a pathogenic MHC class II: Insulin peptide epitope postpone spontaneous autoimmune diabetes. Front Immunol 14, 1207108. 10.3389/fimmu.2023.1207108.

24. Spanier, J.A., Fung, V., Wardell, C.M., Alkhatib, M.H., Chen, Y., Swanson, L.A., Dwyer, A.J., Weno, M.E., Silva, N., Mitchell, J.S., et al. (2023). Tregs with an MHC class II peptide-specific chimeric antigen receptor prevent autoimmune diabetes in mice. J Clin Invest 133. 10.1172/JCI168601.

25. Joffre, O., Santolaria, T., Calise, D., Al Saati, T., Hudrisier, D., Romagnoli, P., and van Meerwijk, J.P. (2008). Prevention of acute and chronic allograft rejection with CD4+CD25+Foxp3+ regulatory T lymphocytes. Nat Med 14, 88–92. 10.1038/nm1688.

26. Tang, Q., Henriksen, K.J., Bi, M., Finger, E.B., Szot, G., Ye, J., Masteller, E.L., McDevitt, H., Bonyhadi, M., and Bluestone, J.A. (2004). In vitro-expanded antigen-specific regulatory T cells suppress autoimmune diabetes. J Exp Med 199, 1455–1465. 10.1084/jem.20040139.

27. Boroughs, A.C., Larson, R.C., Choi, B.D., Bouffard, A.A., Riley, L.S., Schiferle, E., Kulkarni, A.S., Cetrulo, C.L., Ting, D., Blazar, B.R., et al. (2019). Chimeric antigen receptor costimulation domains modulate human regulatory T cell function. JCI Insight 5. 10.1172/jci.insight.126194.

28. Bolivar-Wagers, S., Loschi, M.L., Jin, S., Thangavelu, G., Larson, J.H., McDonald-Hyman, C.S., Aguilar, E.G., Saha, A., Koehn, B.H., Hefazi, M., et al. (2022). Murine CAR19 Tregs suppress acute graft-versus-host disease and maintain graft-versus-tumor responses. JCI Insight 7. 10.1172/jci.insight.160674.

29. Schreeb, K., Culme-Seymour, E., Ridha, E., Dumont, C., Atkinson, G., Hsu, B., and Reinke, P. (2022). Study Design: Human Leukocyte Antigen Class I Molecule A (*)02-Chimeric Antigen Receptor Regulatory T Cells in Renal Transplantation. Kidney Int Rep 7, 1258–1267. 10.1016/j.ekir.2022.03.030.

30. Fritsche, E., Volk, H.D., Reinke, P., and Abou-El-Enein, M. (2020). Toward an Optimized Process for Clinical Manufacturing of CAR-Treg Cell Therapy. Trends Biotechnol 38, 1099–1112. 10.1016/j.tibtech.2019.12.009.

31. Esensten, J.H., Helou, Y.A., Chopra, G., Weiss, A., and Bluestone, J.A. (2016). CD28 Costimulation: From Mechanism to Therapy. Immunity 44, 973–988. 10.1016/j.immuni.2016.04.020.

32. Holst, J., Wang, H., Eder, K.D., Workman, C.J., Boyd, K.L., Baquet, Z., Singh, H., Forbes, K., Chruscinski, A., Smeyne, R., et al. (2008). Scalable signaling mediated by T cell antigen receptor-CD3 ITAMs ensures effective negative selection and prevents autoimmunity. Nat Immunol 9, 658–666. 10.1038/ni.1611.

33. Li, M.O., and Rudensky, A.Y. (2016). T cell receptor signalling in the control of regulatory T cell differentiation and function. Nat Rev Immunol 16, 220–233. 10.1038/nri.2016.26.

34. Yan, D., Farache, J., Mingueneau, M., Mathis, D., and Benoist, C. (2015). Imbalanced signal transduction in regulatory T cells expressing the transcription factor FoxP3. Proc Natl Acad Sci U S A 112, 14942–14947. 10.1073/pnas.1520393112.

35. Crellin, N.K., Garcia, R.V., and Levings, M.K. (2007). Altered activation of AKT is required for the suppressive function of human CD4+CD25+ T regulatory cells. Blood 109, 2014–2022. 10.1182/blood-2006-07-035279.

36. Bloemberg, D., Nguyen, T., MacLean, S., Zafer, A., Gadoury, C., Gurnani, K., Chattopadhyay, A., Ash, J., Lippens, J., Harcus, D., et al. (2020). A High-Throughput Method for Characterizing Novel Chimeric Antigen Receptors in Jurkat Cells. Mol Ther Methods Clin Dev 16, 238–254. 10.1016/j.omtm.2020.01.012.

37. Liu, W., Putnam, A.L., Xu-Yu, Z., Szot, G.L., Lee, M.R., Zhu, S., Gottlieb, P.A., Kapranov, P., Gingeras, T.R., Fazekas de St Groth, B., et al. (2006). CD127 expression inversely correlates with FoxP3 and suppressive function of human CD4+ T reg cells. J Exp Med 203, 1701–1711. 10.1084/jem.20060772.

38. Seddiki, N., Santner-Nanan, B., Martinson, J., Zaunders, J., Sasson, S., Landay, A., Solomon, M., Selby, W., Alexander, S.I., Nanan, R., et al. (2006). Expression of interleukin (IL)-2 and IL-7 receptors discriminates between human regulatory and activated T cells. J Exp Med 203, 1693–1700. 10.1084/jem.20060468.

39. Suhoski, M.M., Golovina, T.N., Aqui, N.A., Tai, V.C., Varela-Rohena, A., Milone, M.C., Carroll, R.G., Riley, J.L., and June, C.H. (2007). Engineering artificial antigen-presenting cells to express a diverse array of co-stimulatory molecules. Mol Ther 15, 981–988. 10.1038/mt.sj.6300134.

40. Ghorashian, S., Kramer, A.M., Onuoha, S., Wright, G., Bartram, J., Richardson, R., Albon, S.J., Casanovas-Company, J., Castro, F., Popova, B., et al. (2019). Enhanced CAR T cell expansion and prolonged persistence in pediatric patients with ALL treated with a low-affinity CD19 CAR. Nat Med 25, 1408–1414. 10.1038/s41591-019-0549-5.

41. Hogquist, K.A., and Jameson, S.C. (2014). The self-obsession of T cells: how TCR signaling thresholds affect fate ‘decisions’ and effector function. Nat Immunol 15, 815–823. 10.1038/ni.2938.

42. Mao, R., Kong, W., and He, Y. (2022). The affinity of antigen-binding domain on the antitumor efficacy of CAR T cells: Moderate is better. Front Immunol 13, 1032403. 10.3389/fimmu.2022.1032403.

43. Abbas, A.K., Trotta, E., D, R.S., Marson, A., and Bluestone, J.A. (2018). Revisiting IL-2: Biology and therapeutic prospects. Sci Immunol 3. 10.1126/sciimmunol.aat1482.

44. Bailey-Bucktrout, S.L., Martinez-Llordella, M., Zhou, X., Anthony, B., Rosenthal, W., Luche, H., Fehling, H.J., and Bluestone, J.A. (2013). Self-antigen-driven activation induces instability of regulatory T cells during an inflammatory autoimmune response. Immunity 39, 949–962. 10.1016/j.immuni.2013.10.016.

45. Overacre-Delgoffe, A.E., Chikina, M., Dadey, R.E., Yano, H., Brunazzi, E.A., Shayan, G., Horne, W., Moskovitz, J.M., Kolls, J.K., Sander, C., et al. (2017). Interferon-gamma Drives T(reg) Fragility to Promote Anti-tumor Immunity. Cell 169, 1130–1141 e1111. 10.1016/j.cell.2017.05.005.

46. Hoffmann, P., Boeld, T.J., Eder, R., Huehn, J., Floess, S., Wieczorek, G., Olek, S., Dietmaier, W., Andreesen, R., and Edinger, M. (2009). Loss of FOXP3 expression in natural human CD4+CD25+ regulatory T cells upon repetitive in vitro stimulation. Eur J Immunol 39, 1088–1097. 10.1002/eji.200838904.

47. Nakagawa, H., Sido, J.M., Reyes, E.E., Kiers, V., Cantor, H., and Kim, H.J. (2016). Instability of Helios-deficient Tregs is associated with conversion to a T-effector phenotype and enhanced antitumor immunity. Proc Natl Acad Sci U S A 113, 6248–6253. 10.1073/pnas.1604765113.

48. Zimmerman, C.M., Robino, R.A., Cochrane, R.W., Dominguez, M.D., and Ferreira, L.M.R. (2024). Redirecting Human Conventional and Regulatory T Cells Using Chimeric Antigen Receptors. Methods Mol Biol 2748, 201–241. 10.1007/978-1-0716-3593-3_15.

49. Collison, L.W., and Vignali, D.A. (2011). In vitro Treg suppression assays. Methods Mol Biol 707, 21–37. 10.1007/978-1-61737-979-6_2.

50. Dawson, N.A.J., Rosado-Sanchez, I., Novakovsky, G.E., Fung, V.C.W., Huang, Q., McIver, E., Sun, G., Gillies, J., Speck, M., Orban, P.C., et al. (2020). Functional effects of chimeric antigen receptor co-receptor signaling domains in human regulatory T cells. Sci Transl Med 12. 10.1126/scitranslmed.aaz3866.

51. Trapani, J.A., and Smyth, M.J. (2002). Functional significance of the perforin/granzyme cell death pathway. Nat Rev Immunol 2, 735–747. 10.1038/nri911.

52. Boissonnas, A., Scholer-Dahirel, A., Simon-Blancal, V., Pace, L., Valet, F., Kissenpfennig, A., Sparwasser, T., Malissen, B., Fetler, L., and Amigorena, S. (2010). Foxp3+ T cells induce perforin-dependent dendritic cell death in tumor-draining lymph nodes. Immunity 32, 266–278. 10.1016/j.immuni.2009.11.015.

53. Ludwig-Portugall, I., Hamilton-Williams, E.E., Gottschalk, C., and Kurts, C. (2008). Cutting edge: CD25+ regulatory T cells prevent expansion and induce apoptosis of B cells specific for tissue autoantigens. J Immunol 181, 4447–4451. 10.4049/jimmunol.181.7.4447.

54. Cao, X., Cai, S.F., Fehniger, T.A., Song, J., Collins, L.I., Piwnica-Worms, D.R., and Ley, T.J. (2007). Granzyme B and perforin are important for regulatory T cell-mediated suppression of tumor clearance. Immunity 27, 635–646. 10.1016/j.immuni.2007.08.014.

55. Grossman, W.J., Verbsky, J.W., Barchet, W., Colonna, M., Atkinson, J.P., and Ley, T.J. (2004). Human T regulatory cells can use the perforin pathway to cause autologous target cell death. Immunity 21, 589–601. 10.1016/j.immuni.2004.09.002.

56. Brinkman, E.K., Chen, T., Amendola, M., and van Steensel, B. (2014). Easy quantitative assessment of genome editing by sequence trace decomposition. Nucleic Acids Res 42, e168 10.1093/nar/gku936.

57. Jennings, E., Elliot, T.A.E., Thawait, N., Kanabar, S., Yam-Puc, J.C., Ono, M., Toellner, K.M., Wraith, D.C., Anderson, G., and Bending, D. (2020). Nr4a1 and Nr4a3 Reporter Mice Are Differentially Sensitive to T Cell Receptor Signal Strength and Duration. Cell Rep 33, 108328. 10.1016/j.celrep.2020.108328.

58. Sundstedt, A., O’Neill, E.J., Nicolson, K.S., and Wraith, D.C. (2003). Role for IL-10 in suppression mediated by peptide-induced regulatory T cells in vivo. J Immunol 170, 1240–1248. 10.4049/jimmunol.170.3.1240.

59. Collison, L.W., Workman, C.J., Kuo, T.T., Boyd, K., Wang, Y., Vignali, K.M., Cross, R., Sehy, D., Blumberg, R.S., and Vignali, D.A. (2007). The inhibitory cytokine IL-35 contributes to regulatory T-cell function. Nature 450, 566–569. 10.1038/nature06306.

60. Kidani, Y., Nogami, W., Yasumizu, Y., Kawashima, A., Tanaka, A., Sonoda, Y., Tona, Y., Nashiki, K., Matsumoto, R., Hagiwara, M., et al. (2022). CCR8-targeted specific depletion of clonally expanded Treg cells in tumor tissues evokes potent tumor immunity with long-lasting memory. Proc Natl Acad Sci U S A 119. 10.1073/pnas.2114282119.

61. Mercer, F., Kozhaya, L., and Unutmaz, D. (2010). Expression and function of TNF and IL-1 receptors on human regulatory T cells. PLoS One 5, e8639. 10.1371/journal.pone.0008639.

62. Subramanian, A., Tamayo, P., Mootha, V.K., Mukherjee, S., Ebert, B.L., Gillette, M.A., Paulovich, A., Pomeroy, S.L., Golub, T.R., Lander, E.S., and Mesirov, J.P. (2005). Gene set enrichment analysis: a knowledge-based approach for interpreting genome-wide expression profiles. Proc Natl Acad Sci U S A 102, 15545–15550. 10.1073/pnas.0506580102.

63. Thornton, A.M., and Shevach, E.M. (2000). Suppressor effector function of CD4+CD25+ immunoregulatory T cells is antigen nonspecific. J Immunol 164, 183–190. 10.4049/jimmunol.164.1.183.

64. Elgueta, R., Benson, M.J., de Vries, V.C., Wasiuk, A., Guo, Y., and Noelle, R.J. (2009). Molecular mechanism and function of CD40/CD40L engagement in the immune system. Immunol Rev 229, 152–172. 10.1111/j.1600-065X.2009.00782.x.

65. Gu, J., Ni, X., Pan, X., Lu, H., Lu, Y., Zhao, J., Guo Zheng, S., Hippen, K.L., Wang, X., and Lu, L. (2017). Human CD39(hi) regulatory T cells present stronger stability and function under inflammatory conditions. Cell Mol Immunol 14, 521–528. 10.1038/cmi.2016.30.

66. Harshe, R.P., Xie, A., Vuerich, M., Frank, L.A., Gromova, B., Zhang, H., Robles, R.J., Mukherjee, S., Csizmadia, E., Kokkotou, E., et al. (2020). Endogenous antisense RNA curbs CD39 expression in Crohn’s disease. Nat Commun 11, 5894. 10.1038/s41467-020-19692-y.

67. Bin Dhuban, K., d’Hennezel, E., Nashi, E., Bar-Or, A., Rieder, S., Shevach, E.M., Nagata, S., and Piccirillo, C.A. (2015). Coexpression of TIGIT and FCRL3 identifies Helios+ human memory regulatory T cells. J Immunol 194, 3687–3696. 10.4049/jimmunol.1401803.

68. Joller, N., Lozano, E., Burkett, P.R., Patel, B., Xiao, S., Zhu, C., Xia, J., Tan, T.G., Sefik, E., Yajnik, V., et al. (2014). Treg cells expressing the coinhibitory molecule TIGIT selectively inhibit proinflammatory Th1 and Th17 cell responses. Immunity 40, 569–581. 10.1016/j.immuni.2014.02.012.

69. Lee, D.J. (2020). The relationship between TIGIT(+) regulatory T cells and autoimmune disease. Int Immunopharmacol 83, 106378. 10.1016/j.intimp.2020.106378.

70. Hu, X., and Ivashkiv, L.B. (2009). Cross-regulation of signaling pathways by interferon-gamma: implications for immune responses and autoimmune diseases. Immunity 31, 539–550. 10.1016/j.immuni.2009.09.002.

71. DuPage, M., and Bluestone, J.A. (2016). Harnessing the plasticity of CD4(+) T cells to treat immune-mediated disease. Nat Rev Immunol 16, 149–163. 10.1038/nri.2015.18.

72. Schmid, D.A., Irving, M.B., Posevitz, V., Hebeisen, M., Posevitz-Fejfar, A., Sarria, J.C., Gomez-Eerland, R., Thome, M., Schumacher, T.N., Romero, P., et al. (2010). Evidence for a TCR affinity threshold delimiting maximal CD8 T cell function. J Immunol 184, 4936–4946. 10.4049/jimmunol.1000173.

73. Chen, L., and Flies, D.B. (2013). Molecular mechanisms of T cell co-stimulation and co-inhibition. Nat Rev Immunol 13, 227–242 10.1038/nri3405.

74. Salter, A.I., Ivey, R.G., Kennedy, J.J., Voillet, V., Rajan, A., Alderman, E.J., Voytovich, U.J., Lin, C., Sommermeyer, D., Liu, L., et al. (2018). Phosphoproteomic analysis of chimeric antigen receptor signaling reveals kinetic and quantitative differences that affect cell function. Sci Signal 11. 10.1126/scisignal.aat6753.

75. Fisher, J.D., Zhang, W., Balmert, S.C., Aral, A.M., Acharya, A.P., Kulahci, Y., Li, J., Turnquist, H.R., Thomson, A.W., Solari, M.G., et al. (2020). In situ recruitment of regulatory T cells promotes donor-specific tolerance in vascularized composite allotransplantation. Sci Adv 6, eaax8429. 10.1126/sciadv.aax8429.

76. Boroughs, A.C., Larson, R.C., Marjanovic, N.D., Gosik, K., Castano, A.P., Porter, C.B.M., Lorrey, S.J., Ashenberg, O., Jerby, L., Hofree, M., et al. (2020). A Distinct Transcriptional Program in Human CAR T Cells Bearing the 4-1BB Signaling Domain Revealed by scRNA-Seq. Mol Ther 28, 2577–2592. 10.1016/j.ymthe.2020.07.023.

77. Salter, A.I., Rajan, A., Kennedy, J.J., Ivey, R.G., Shelby, S.A., Leung, I., Templeton, M.L., Muhunthan, V., Voillet, V., Sommermeyer, D., et al. (2021). Comparative analysis of TCR and CAR signaling informs CAR designs with superior antigen sensitivity and in vivo function. Sci Signal 14. 10.1126/scisignal.abe2606.

78. Davenport, A.J., Cross, R.S., Watson, K.A., Liao, Y., Shi, W., Prince, H.M., Beavis, P.A., Trapani, J.A., Kershaw, M.H., Ritchie, D.S., et al. (2018). Chimeric antigen receptor T cells form nonclassical and potent immune synapses driving rapid cytotoxicity. Proc Natl Acad Sci U S A 115, E2068–E2076. 10.1073/pnas.1716266115.

79. Pandiyan, P., Zheng, L., Ishihara, S., Reed, J., and Lenardo, M.J. (2007). CD4+CD25+Foxp3+ regulatory T cells induce cytokine deprivation-mediated apoptosis of effector CD4+ T cells. Nat Immunol 8, 1353–1362. 10.1038/ni1536.

80. Dalton, D.K., Pitts-Meek, S., Keshav, S., Figari, I.S., Bradley, A., and Stewart, T.A. (1993). Multiple defects of immune cell function in mice with disrupted interferon-gamma genes. Science 259, 1739–1742. 10.1126/science.8456300.

81. Marson, A., Kretschmer, K., Frampton, G.M., Jacobsen, E.S., Polansky, J.K., MacIsaac, K.D., Levine, S.S., Fraenkel, E., von Boehmer, H., and Young, R.A. (2007). Foxp3 occupancy and regulation of key target genes during T-cell stimulation. Nature 445, 931–935. 10.1038/nature05478.

82. Esposito, M., Ruffini, F., Bergami, A., Garzetti, L., Borsellino, G., Battistini, L., Martino, G., and Furlan, R. (2010). IL-17- and IFN-gamma-secreting Foxp3+ T cells infiltrate the target tissue in experimental autoimmunity. J Immunol 185, 7467–7473. 10.4049/jimmunol.1001519.

83. Duhen, T., Duhen, R., Lanzavecchia, A., Sallusto, F., and Campbell, D.J. (2012). Functionally distinct subsets of human FOXP3+ Treg cells that phenotypically mirror effector Th cells. Blood 119, 4430–4440. 10.1182/blood-2011-11-392324.

84. Schoenbrunn, A., Frentsch, M., Kohler, S., Keye, J., Dooms, H., Moewes, B., Dong, J., Loddenkemper, C., Sieper, J., Wu, P., et al. (2012). A converse 4-1BB and CD40 ligand expression pattern delineates activated regulatory T cells (Treg) and conventional T cells enabling direct isolation of alloantigen-reactive natural Foxp3+ Treg. J Immunol 189, 5985–5994. 10.4049/jimmunol.1201090.

85. Nowak, A., Lock, D., Bacher, P., Hohnstein, T., Vogt, K., Gottfreund, J., Giehr, P., Polansky, J.K., Sawitzki, B., Kaiser, A., et al. (2018). CD137+CD154-Expression As a Regulatory T Cell (Treg)-Specific Activation Signature for Identification and Sorting of Stable Human Tregs from In Vitro Expansion Cultures. Front Immunol 9, 199. 10.3389/fimmu.2018.00199.

86. Shen, X., Wang, Y., Gao, F., Ren, F., Busuttil, R.W., Kupiec-Weglinski, J.W., and Zhai, Y. (2009). CD4 T cells promote tissue inflammation via CD40 signaling without de novo activation in a murine model of liver ischemia/reperfusion injury. Hepatology 50, 1537–1546. 10.1002/hep.23153.

87. Majzner, R.G., Rietberg, S.P., Sotillo, E., Dong, R., Vachharajani, V.T., Labanieh, L., Myklebust, J.H., Kadapakkam, M., Weber, E.W., Tousley, A.M., et al. (2020). Tuning the Antigen Density Requirement for CAR T-cell Activity. Cancer Discov 10, 702–723. 10.1158/2159-8290.CD-19-0945.

88. Muller, Y.D., Nguyen, D.P., Ferreira, L.M.R., Ho, P., Raffin, C., Valencia, R.V.B., Congrave-Wilson, Z., Roth, T.L., Eyquem, J., Van Gool, F., et al. (2021). The CD28-Transmembrane Domain Mediates Chimeric Antigen Receptor Heterodimerization With CD28. Front Immunol 12, 639818. 10.3389/fimmu.2021.639818.

89. Ferreira, L.M.R., Kaul, A.M., Guerrero-Moreno, R., Fontenot, J.D., Bluestone, J.A., and Tang, Q. (2018). Tailoring a new generation of chimeric antigen receptors for regulatory T cells. J Immunol 200 ((*1_Supplement*)), 176.175.

90. Fung, V.C.W., Rosado-Sanchez, I., and Levings, M.K. (2021). Transduction of Human T Cell Subsets with Lentivirus. Methods Mol Biol 2285, 227–254. 10.1007/978-1-0716-1311-5_19.

91. Dobin, A., Davis, C.A., Schlesinger, F., Drenkow, J., Zaleski, C., Jha, S., Batut, P., Chaisson, M., and Gingeras, T.R. (2013). STAR: ultrafast universal RNA-seq aligner. Bioinformatics 29, 15–21. 10.1093/bioinformatics/bts635.

92. Liao, Y., Smyth, G.K., and Shi, W. (2014). featureCounts: an efficient general purpose program for assigning sequence reads to genomic features. Bioinformatics 30, 923–930. 10.1093/bioinformatics/btt656.

93. Love, M.I., Huber, W., and Anders, S. (2014). Moderated estimation of fold change and dispersion for RNA-seq data with DESeq2. Genome Biol 15, 550. 10.1186/s13059-014-0550-8.

94. Ge, S.X., Son, E.W., and Yao, R. (2018). iDEP: an integrated web application for differential expression and pathway analysis of RNA-Seq data. BMC Bioinformatics 19, 534. 10.1186/s12859-018-2486-6.

