## Supplemental Table 9 for "High affinity chimeric antigen receptor signaling induces an inflammatory program in human regulatory T cells"

| **Antigen** | **Clone** | **Fluorophore** | **Dilution** | **Vendor** | **Catalog #** |
| --- | --- | --- | --- | --- | --- |
| CD4 | SK3 | FITC | 1:100 | BioLegend | 980802 |
| CD4 | SK3 | Pacific Blue | 1:100 | BioLegend | 344619 |
| CD4 | SK3 | Alexa Fluor 700 | 1:100 | BioLegend | 344621 |
| MYC-tag | 9B11 | Alexa Fluor 647 | 1:100 | Cell Signaling Technologies | 2233S |
| CD8 | SK1 | PE | 1:100 | BioLegend | 344706 |
| CD8 | SK1 | PerCP | 1:100 | BioLegend | 344707 |
| CD25 | BC96 | APC | 1:100 | BioLegend | 302610 |
| CD71 | CY1G4 | PE | 1:100 | BioLegend | 334105 |
| CD80 | 2D10 | APC | 1:100 | BioLegend | 305220 |
| CD83 | HB15e | Brilliant Violet 421 | 1:100 | BioLegend | 305324 |
| CD86 | IT2.2 | PE | 1:100 | BioLegend | 305406 |
| CD127 | A019D5 | PE | 1:100 | BioLegend | 351304 |
| CellTrace  Violet | N/A | CellTrace  Violet | 1:1000 | ThermoFisher | [C34571](https://www.thermofisher.com/order/catalog/product/C34571) |
| CellTrace Far Red | N/A | CellTrace Far Red | 1:350 | ThermoFisher | C34572 |
| FOXP3 | PCH101 | eFluor 450 | 1:50 | eBioscience | 48-4776-42 |
| FOXP3 | PCH101 | PE/Cy5.5 | 1:50 | ThermoFisher | 35-4776-42 |
| IFNG | 4S.B3 | BV510 | 1:50 | Biolegend | 502543 |
| IL2 | MQ1-17H12 | Alexa Fluor 647 | 1:50 | Biolegend | 500315 |
| CD154 | 24-31 | APC | 1:100 | Biolegend | 310809 |
| TIGIT | A15153G | Brilliant Violet 785 | 1:100 | Biolegend | 372735 |
| Ghost | N/A | Red 780 | 1:500 | Tonbo Biosciences | 13-0865-T500 |
| HELIOS | 22F6 | PE | 1:50 | BioLegend | 137216 |
| Live-or-Dye | N/A | 594/614 | 1:500 | Biotium | 32006 |
